# Simultaneous acclimation to nitrogen and iron scarcity in open ocean cyanobacteria revealed by sparse tensor decomposition of metatranscriptomes

**DOI:** 10.1101/2024.07.15.603627

**Authors:** Stephen Blaskowski, Marie Roald, Paul M. Berube, Rogier Braakman, E. Virginia Armbrust

**Affiliations:** Molecular Engineering and Sciences Institute, University of Washington, Seattle, Washington, USA; School of Oceanography, University of Washington, Seattle, Washington, USA; Simula Metropolitan Center for Digital Engineering, Oslo, Oslo, Norway; Faculty of Technology, Art and Design, Oslo Metropolitan University, Oslo, Oslo, Norway; Department of Civil and Environmental Engineering, Massachusetts Institute of Technology, Cambridge, Massachusetts, USA; Department of Earth, Atmospheric and Planetary Sciences, Massachusetts Institute of Technology, Cambridge, Massachusetts, USA

## Abstract

Microbes respond to changes in their environment by adapting their physiology through coordinated adjustments to the expression levels of functionally related genes. To detect these shifts in situ, we developed a sparse tensor decomposition method that derives gene co-expression patterns from inherently complex whole community RNA-sequencing data. Application of the method to metatranscriptomes of the abundant marine cyanobacteria *Prochlorococcus* and *Synechococcus* identified responses to scarcity of two essential nutrients, nitrogen and iron, including increased transporter expression, restructured photosynthesis and carbon metabolism, and mitigation of oxidative stress. Further, expression profiles of the identified gene clusters suggest that both cyanobacteria populations experience simultaneous nitrogen and iron stresses in a transition zone between North Pacific oceanic gyres. The results demonstrate the power of our approach to infer organism responses to environmental pressures, hypothesize functions of uncharacterized genes, and extrapolate ramifications for biogeochemical cycles in a changing ecosystem.

**Teaser:** New analytical approach reveals shifts in gene expression that may help cyanobacteria cope with environmental stressors.

## Introduction

Microbial communities play essential roles in every Earth biome and drive global biogeochemical cycles through their collective metabolic activity (*1*). In open ocean ecosystems, the abundant cyanobacteria *Prochlorococcus* and *Synechococcus* are keystone photosynthetic microbes that account for an estimated *⇠* 20 *-* 25% of total marine primary production (*2, 3*). Although descended from a common ancestor, these two genera encompass a diversity of genetically and phenotypically differentiated subpopulations (*4*). *Prochlorococcus* numerically dominates in the nutrient-poor tropical and subtropical open ocean waters that span from about 40°S to 40°N, whereas *Synechococcus* thrives across a greater range of habitats and is more abundant in colder, more nutrient-rich regions (*2, 3*). Within each genus, genetically differentiated clades are further adapted to occupy distinct ecological niches, such as high-light adapted *Prochlorococcus* clades encountered near the surface, and low-light adapted *Prochlorococcus* clades generally found in deeper waters (*5*). This ecological partitioning is driven by the functional diversity of an extensive pangenome (*6*). *Prochlorococcus* and *Synechococcus* genomes share a core of about 1,000 – 1,500 genes, and a suite of accessory genes enable different clades and strains to adapt to different environmental conditions (*5, 7*). Cultured isolates do not yet represent the full breadth of this sequence diversity, and a majority of the cyanobacterial pangenome remains functionally uncharacterized – part of the “functional dark matter” that constitutes a majority of gene families in the overall global microbiome (*8*). Moreover, metabolism emerges not from the activity of independent genes, but rather from multiple gene products acting in concert. Characterizing these functional gene networks in marine cyanobacteria is crucial for clarifying the metabolic links between microbial genes and the ecosystem-scale processes they drive.

In recent decades, omics technologies have proven to be powerful tools for studying microbial communities, generating comprehensive inventories of the DNA, RNA, proteins, and metabolites present in a microbiome (*9*). Metatranscriptomic datasets are particularly information-rich because they catalog both the sequence of an RNA transcript, imprinted with the identity of the originating gene and organism, as well as the transcript’s quantity, a composite reflection of organism abundance and gene expression. Thus, metatranscriptomes capture a snapshot of the in situ physiological dynamics of a microbiome. Commonly researchers infer these dynamics from metatranscriptomic data by comparing between sampling conditions the expression levels of genes involved in metabolic pathways of interest; however this approach is dependent on prior knowledge of the genes and pathways in question. Unsupervised clustering analysis provides an alternate avenue, independent of prior knowledge, which instead identifies groups of transcripts with similar abundance profiles across sampling conditions, aiming to uncover cohorts of co-expressed genes. These gene co-expression clusters reflect the modular organization of metabolic pathways and gene regulatory networks, and allow the functions of un-annotated genes to be inferred based on their association with better-characterized genes. Currently, the accumulation of metatranscriptomic data exceeds the advancement analytical frameworks, and co-expression clustering represents a critical part of this gap.

Several co-expression clustering methods have been designed specifically for metatranscriptomic datasets (*10–12*), and additional studies have relied on methods originally designed for analyzing single-organism RNA sequencing data (*13*). None of these approaches consider taxonomic information in the cluster detection algorithm, and resulting clusters can range from hundreds to thousands of genes, limiting interpretive power. Single-organism approaches are in turn ill-equipped to handle the idiosyncratic properties of metatranscriptomic data. Typical metatranscriptomic datasets are characterized by high dimensionality, variable community composition, overdispersion (variance increases exponentially with increasing mean), technical noise, and pervasive zero values, each presenting an obstacle to effective analysis (*14*). And yet, the complexity of metatranscriptomic data also provides an opportunity. Metatranscriptomic datasets have a multiway structure, in which each transcript abundance data point can be considered indexed by three variables: gene, taxon, and sample, with the latter serving as a proxy for the environmental context of expression. Analysis that effectively models this multiway structure can produce more robust inferences about the patterns of gene expression that give rise to metatranscriptomes, elucidating previously unrecognized microbiome dynamics.

Here we introduce Barnacle, a pattern discovery method developed to identify interpretable clusters of genes co-expressed across metatranscriptomes. Barnacle leverages the inherent multiway structure of metatranscriptomic data in combination with sparsity and non-negativity constraints to identify gene co-expression clusters that reflect the physiological states of organisms interacting in an environment. The foundation of the approach is CANDECOMP/PARAFAC (CP) tensor decomposition, a technique that models a multiway dataset as a sum of its constituent signals (for a review of tensor decomposition see Kolda and Bader, 2009 (*15*)). Component models, which include CP tensor decomposition, stand out among gene clustering tech-niques for their accuracy and robustness to noise, as well as their capacity to accommodate gene membership in more than one cluster, a property reflective of the structure of metabolic and regulatory networks (*16*). Tensor decomposition methods have been developed for analyzing human gene expression datasets, including two studies that informed the formulation of model constraints used in Barnacle: the SDA algorithm that incorporates sparsity constraints using a Bayesian framework (*17*), and the MultiCluster method that employs partial non-negativity constraints (*18*). Our approach represents a novel application of tensor decomposition to metatranscriptomic data, developed in conjunction with innovative procedures for parameter selection and bootstrapping, that together produce robust, interpretable inferences.

In this study we describe construction of the sparse tensor decomposition model and validate its functionality using simulated data. We then turn Barnacle’s capabilities to the analysis of 222 marine metatranscriptomes collected during three cruises in the North Pacific Ocean, which transited along the 158th meridian west (fig. S1), sampling from the nitrogen-limited subtropical gyre to the southernmost edge of the iron-limited subarctic gyre (*19*). The transition zone between the two gyres is a dynamic interface in which elevated phytoplankton productivity supports a high diversity of marine life, including albacore tuna and loggerhead turtles that follow the front as a migratory route (*20*). We focused our analysis on the cyanobacterial communities that form the foundation of these ecosystems and uncovered gene clusters that underlie acclimation of *Prochlorococcus* and *Synechococcus* subpopulations to shifting environmental pressures. This work demonstrates the power of unsupervised signal discovery techniques to draw from meta-omics data explanatory connections between the molecular functions of individual genes, the cellular programs these genes drive, and the ecosystem scale processes that emerge from the collective metabolic activities of interacting microbes.

## Results

We developed a sparse tensor decomposition model, Barnacle, to detect co-expressed gene modules in metatranscriptomes of marine microbiomes. Our modeling framework represents the multiway structure of metatranscriptomic data as a three-way gene *⇥* taxon *⇥* sample tensor of transcript abundance values (see Materials and Methods for complete model description). The model approximates this input data tensor as a sum of components (fig. S2A), each of which represents a pattern of transcript abundance specific to particular genes, taxa, and sampling conditions (fig. S2B). Two key parameters must be tuned during model fitting: *R* and *.λ* (see Table 1 for symbol definitions). Tensor rank (*R*) dictates the number of components used to model the data. The number of genes included in each component is indirectly controlled by *.λ*, a sparsity coefficient that constrains the number of non-zero gene weights in the model. In each component, the genes corresponding to non-zero weights comprise a gene cluster (fig. S2C), and the taxon and sample weights indicate the relative expression of the cluster in each taxon and sample in the dataset. As a result, components can be interpreted as reflecting the physiological states of the populations modeled, generated by defined programs of coordinated gene expression.

**Table 1.** Main symbols used in the text.

| Symbol | Definition |
| --- | --- |
| $R$ | rank (number of components) |
| $\lambda$ | sparsity coefficient |
| $\mathcal{Y}$ | data tensor (gene × taxon × sample) |
| $\mathbf{G}$ | gene component matrix |
| $\mathbf{T}$ | taxon component matrix |
| $\mathbf{S}$ | sample component matrix |
| $\mathbf{g}_r$ | component $r$ of matrix $\mathbf{G}$ (vector) |
| $Y_{(\mathbf{G})}$ | gene-mode unfolding of tensor $\mathcal{Y}$ (matrix) |
| $\ \cdot\ _F$ | Frobenius norm |
| $\circ$ | Outer product |
| $\odot$ | Khatri-Rao product (column-wise Kronecker product) |

### Model validation with simulated data

Model performance was evaluated using simulated data tensors, each constructed as the sum of a specified number of sparse components, combined with a Gaussian noise tensor. We measured performance on the basis of five metrics (Table 2): the sum of squared errors (SSE) to evaluate overall model fit, factor match score (FMS) to compare model components against ground truth components used to generate the simulation, and precision and recall metrics (*16*) to assess the similarity of model-derived gene sets to those derived from simulation components. Additionally, we calculated the harmonic mean of the precision and recall scores to produce the F1 score, providing a balanced measure of cluster accuracy.

**Table 2.**
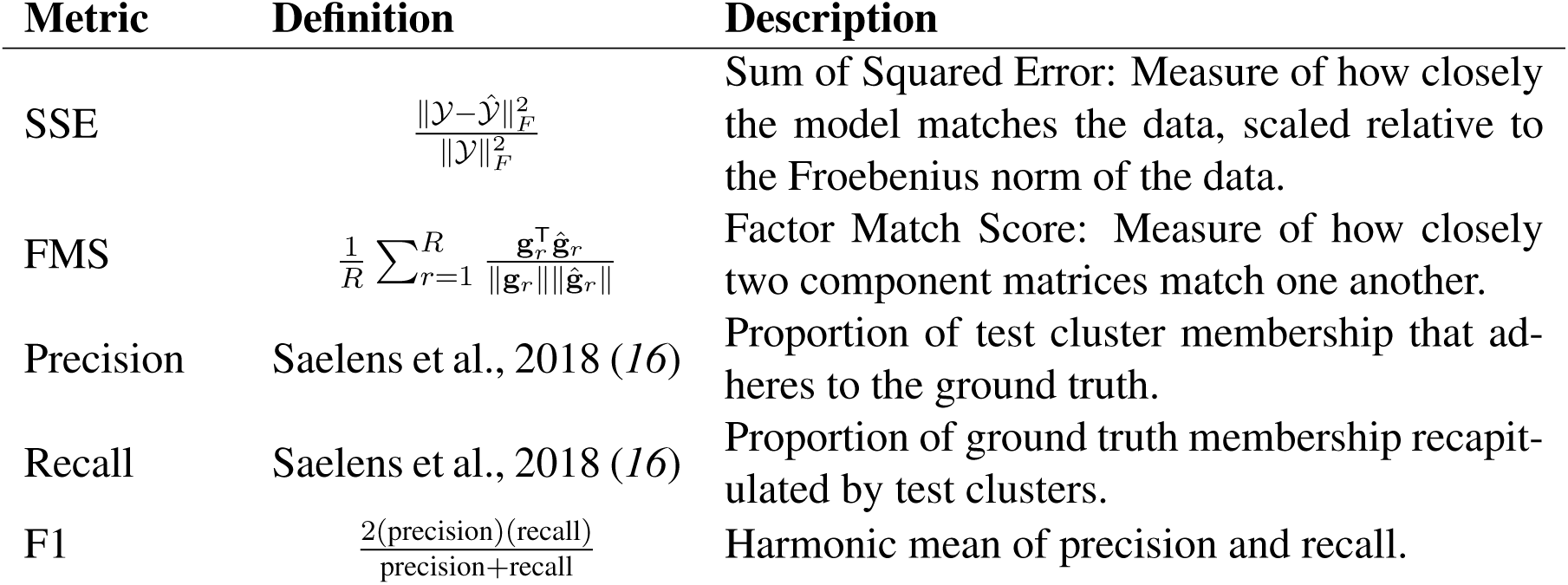
Metrics used to evaluate model performance.

The results of a representative simulation experiment (Fig. 1) illustrate use of these metrics to evaluate model performance. In the experiment shown, we generated a simulated gene *⇥* taxon *⇥* sample tensor of shape 50 *⇥* 20 *⇥* 30 using 8 components with an average gene cluster size of 20 (Fig. 1D, left side) and a noise-to-signal ratio of 1. A series of models parameterized with different numbers of components (*R*) and sparsity coefficients (*.λ*) were fit to the simulated data. Across a range of *.λ* values, the model with the lowest SSE (comparing model to noiseless simulation) corresponded to *R* = 8 (Fig. 1A), indicating that the most accurate model fit was achieved when the number of model components matched the true number of components used to generate the simulation. Given *R* = 8, we evaluated the effect of sparsity coefficient (*.λ*) on model fit (SSE) and the accuracy of model components (FMS) and found that *.λ* = 0.8 corresponded to both the minimum SSE and the maximum FMS (Fig. 1B). However, the maximum F1 score, indicating the best accuracy of model-derived gene clusters, corresponded to *.λ* = 1.4 (Fig. 1C). This value of *.λ* corresponded to the inflection points of the SSE and FMS curves rather than their respective minimum and maximum. The results also highlighted the inherent trade off between precision and recall: gene cluster recall improved at the expense of precision below *.λ* = 1.4, and precision improved at the expense of recall above *.λ* = 1.4 (Fig. 1C). Importantly, the component matrices of the model parameterized with *R* = 8 (corresponding to minimum SSE), and *.λ* = 1.4, (corresponding to maximum F1), closely resembled the ground truth components used to generate the simulation (Fig. 1D).

**Fig. 1.**
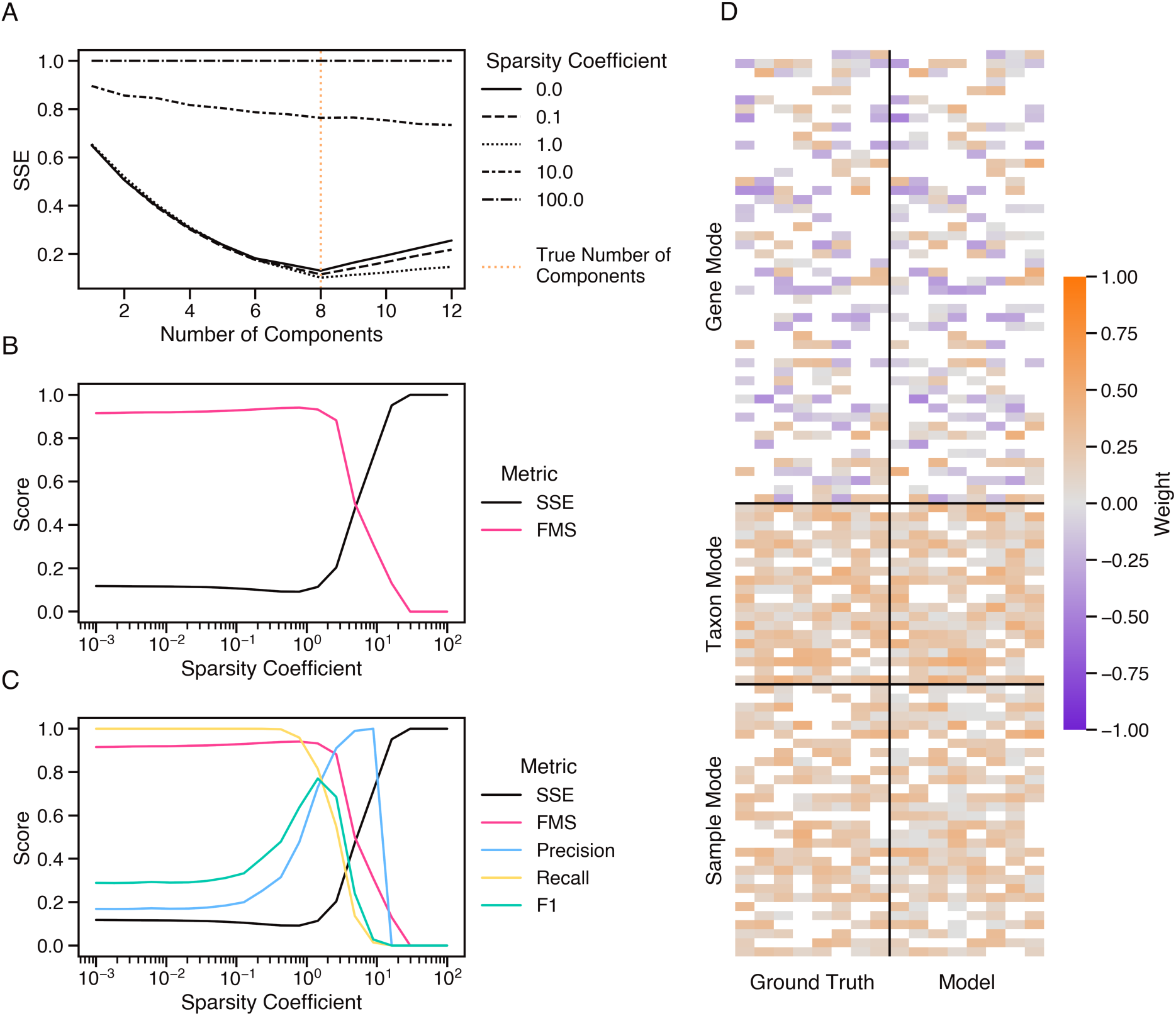
Example model evaluation with noisy simulated data tensor. (A) Changes in the relative sum of squared errors (SSE) resulting from parameterization with different numbers of components (*R*) and sparsity coefficients (*.λ*). Vertical orange line indicates the number of components used to generate the simulation. Simulated noise level was scaled to a 1:1 noise-to-signal ratio. (B) Changes in the factor match score (FMS) and SSE resulting from parameterization with different values of *.λ* (number of components fixed at *R* = 8). (C) Changes in the precision, recall, and F1 score of gene clusters derived from models parameterized with different *.λ* values (fixed *R* = 8), plotted alongside SSE and FMS for ease of comparison. (D) Weights of a model parameterized with *R* = 8 and *.λ* = 1.4 (right), shown in comparison to the components used to construct the simulation (left). Zero-valued weights are indicated by blank spaces.

Using these metrics, we assessed generalized model performance on 100 simulated tensors that collectively represented a range of tensor shapes, ranks, sparsity patterns, and noise-to-signal ratios ranging from 0.1 to 10 (Fig. 2). The sparse tensor decomposition model was fit to each simulation using optimal parameters. The *R* parameter was set to the true number of components used to generate each simulation. The *.λ* parameter does not correspond to any value used to generate simulations, so the optimal sparsity coefficient was estimated as the value of *.λ* that yielded the maximum F1 score, which generally increased with the noise level and fraction of zero values of the simulation (fig. S3). Model fit, as measured by SSE, closely tracked simulated noise level (Fig. 2A), indicating that the model represented the variation attributable to signal and did not over-fit to noise. Model components matched the component matrices used to generate simulations almost identically up to a noise-to-signal ratio of 1, above which FMS decreased to an average of about 0.6 at a noise-to-signal ratio of 10 (Fig. 2B). The precision and recall of model-derived gene clusters were similarly high below a noise-to-signal ratio of 1 and exhibited a more gradual decline at higher noise levels compared to FMS, with average precision around 0.7 and average recall around 0.9 at a noise-to-signal ratio of 10 (Fig. 2, C and D). F1 scores reflected a balance between precision and recall with an average around 0.7 at the highest noise levels tested (Fig. 2E). Overall these results indicate that the sparsity constraint is an effective counterbalance to noise, allowing the model to faithfully recover signal even in high noise datasets, and that this performance does not appear affected by rank, tensor shape, or the proportion of zero values.

**Fig. 2.**
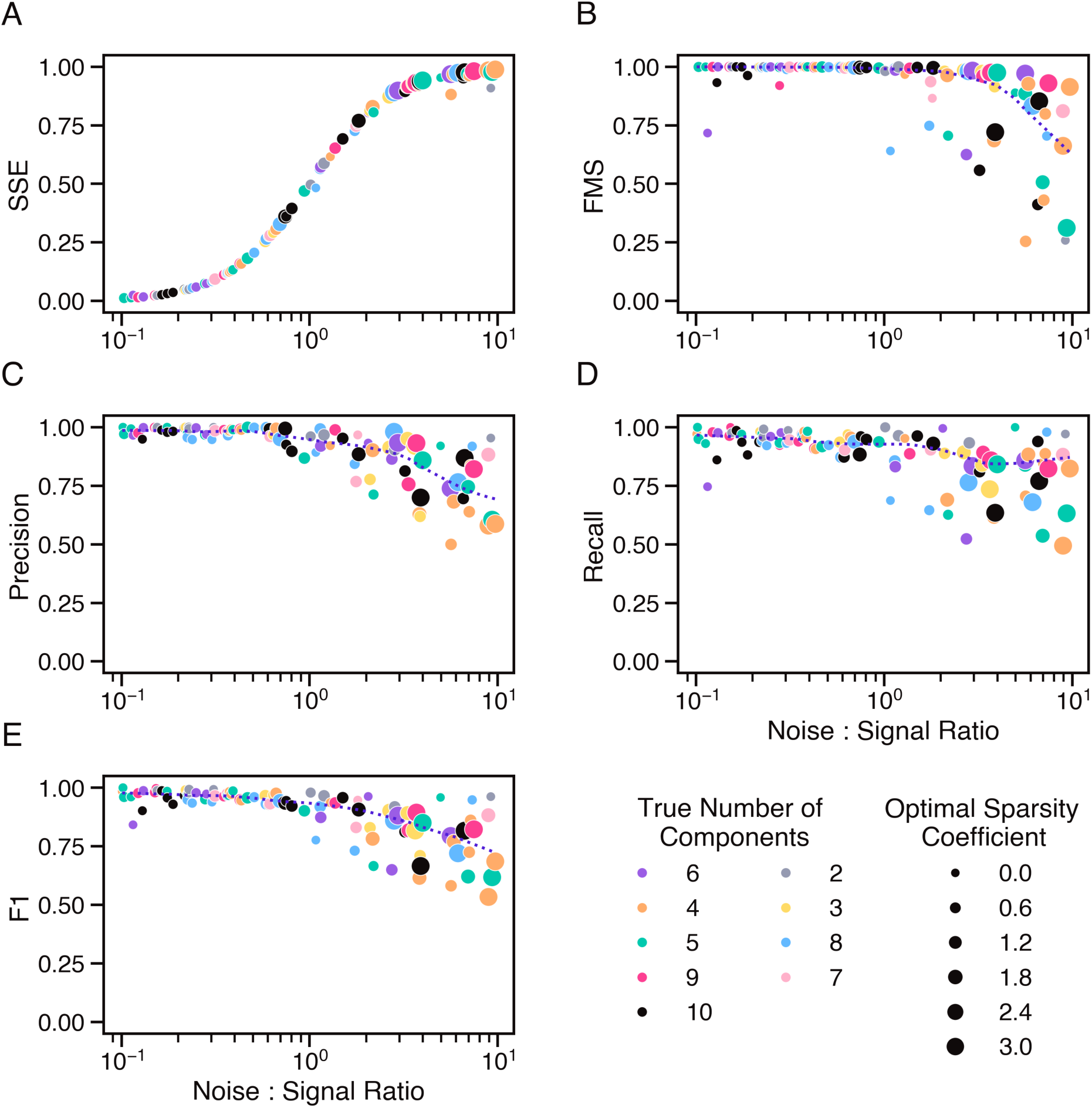
Generalized model performance on 100 noisy simulated data tensors. Performance of optimally-parameterized models of 100 simulated data tensors, evaluated in reference to simulation ground truth and plotted as a function of simulated noise-to-signal ratio. Optimal *R* parameter, indicated by marker color, was set equal to the true number of components used to generate each simulation, and optimal sparsity coefficient, indicated by marker size, was determined by maximum F1 score. (A) Overall model fit as measured by sum of squared errors (SSE). (B) Accuracy of component matrices as measured by factor match score (FMS). Accuracy of gene clusters resulting from model gene components, measured in terms of (C) precision, (D) recall, and (E) F1 score. Dotted lines indicate trends, as determined by locally weighted scatter plot smoothing (LOWESS) with a bandwidth of 0.3.

### Parameter selection

Optimal model parameters are rarely known a priori and instead must be inferred from the data. We therefore developed a cross-validated grid search strategy that leverages sample replicates to select *R* and *.λ* parameters that result in the best fit model (see Materials and Methods). In brief, for each unique set of parameters, we fit a model to a subset of the data encompassing one replicate from each sampling condition, and then calculated cross-validated SSE and FMS scores using the held out replicates. We selected the *R* value of best fit based on the minimum cross-validated SSE. Fixing the *R* parameter, we then selected the best fit sparsity coefficient as the maximum *.λ* value at which the cross-validated FMS remained within one standard error of the maximum FMS, a variation on the 1SE rule for parsimonious sparse model selection (*21*). We assessed the effectiveness of this strategy using the 100 simulated tensors, and generally found that the procedure selected parameters that approached optimal values up to a noise level ten times that of the signal (see Supplementary Text and fig. S4). Even in cases where the procedure led to selecting suboptimal parameters, this generally had a negligible effect on model performance, although significantly undershooting the optimal sparsity coefficient incurred a decrease in cluster precision whereas overshooting incurred a decrease in recall (fig. S5). Although optimal parameters will yield the most accurate model, given a choice between more or less sparsity, these results indicate that erring on the side of greater sparsity will decrease invalid inferences based on false positives.

### Model fitting to metatranscriptomic data

After evaluating model performance on simulated data, we employed Barnacle to identify coexpressed gene clusters in marine metatranscriptomes, in an attempt to deepen our understanding of ecosystem dynamics in open ocean cyanobacteria populations. To this end, we compiled a dataset of 222 metatranscriptomes collected across different years and locations in the North Pacific Ocean (fig. S1). We mapped the metatranscriptome sequencing reads against a database of 681 *Prochlorococcus* and *Synechococcus* reference genomes (*22*) to determine the abundances of transcripts originating from these species. Sequencing read counts were then aggregated by Cyanobacterial Clusters of Orthologous Groups of proteins (CyCOGs) and taxonomic clade. Clade aggregated read counts revealed a latitudinal shift in cyanobacterial community composition that was relatively consistent between cruise years. The *Prochlorococcus* community was dominated by the HLII clade within the North Pacific Subtropical Gyre (NPSG), shifted to HLI dominance in the transition zone, and gave way to a smaller community primarily consisting of LLI and LLVII clades near the southern subarctic gyre (fig. S6). While the *Synechococcus* community showed more variability, predominate trends revealed the dominance of clade II in the NPSG, a rise in the community share of CRD1 and CRD2 in the transition zone, and the prevalence of clades I and IV near the subarctic gyre, as well as some detection of subclusters 5.2 and 5.3 at the northernmost extent of the transects (fig. S7). These observed shifts in community composition underscore the dynamic nature of North Pacific ecosystems, and highlight a persistent challenge in the analysis of environmental metatranscriptomes: disentangling changes in gene expression from changes in organism abundance.

To focus our analysis on patterns of differential gene expression and to adjust for differences between samples in community composition and sequencing depth, we normalized transcript abundance counts by clade. In both the *Prochlorococcus* and *Synechococcus* datasets, gene abundance profiles exhibited an overdispersed mean-variance relationship, indicative of a negative binomial distribution (fig. S8). We therefore normalized read counts using the variance stabilizing transform (vst) which employs a generalized linear model with negative binomial variance to stabilize overdispersion in count matrices (*23, 24*). Normalization with vst produced residual transcript abundance values, which quantify the degree and direction to which each transcript abundance diverged from expectation in each sample, given the distribution of abundance counts across all samples. These values can be described as transcription anomalies, which we interpret as instance of up-regulated or down-regulated gene expression. Genes that do not significantly vary between sampling conditions, after accounting for changes in organism abundance, will result in residual transcript abundance values near zero. Normalization successfully decoupled variance from mean, and the resulting values exhibited a similar range between clades (fig. S9). We then arranged the normalized residual transcript abundance data into two tensorized datasets by aligning CyCOGs across clades. One tensor consisted of *Prochlorococcus* clades HLI, HLII, and LLI. The second tensor consisted of *Synechococcus* clades I, II, III, IV, CRD1, CRD2, and UC-A. *Prochlorococcus* clade LLVII and *Synechococcus* subclusters 5.2 and 5.3 were excluded from our analysis despite their apparent prevalence in some samples (figs. S6 and S7) because those transcript bins mapped to a low proportion of the clade pangenome, indicating a poor match between reference genomes and the sequenced metatranscriptomes.

We applied Barnacle to the normalized *Prochlorococcus* and *Synechococcus* residual transcript abundance tensors to identify clusters of CyCOGs with correlated patterns of anomalous expression, attributable to particular samples and clades. Best fit model parameters were identified using the cross-validated grid search strategy in combination with bootstrapping (fig. S10). Bootstrapping allowed use of all samples for model fitting despite discrepancies in the number of replicates available for each sampling condition (see Materials and Methods and fig. S1B). Among models fit without any applied sparsity penalty (*.λ*=0.0), the minimum cross-validated SSE corresponded to an *R* of 15 components in both the *Prochlorococcus* and *Synechococcus* datasets (fig. S10, A and B). We selected the number of components based on the models with *.λ*=0.0 as opposed to the overall minimum in SSE because among models fit with a sparsity coefficient of *.λ*=10.0, SSE continually decreased with increasing *R* up to a value of *R* = 1600, a point at which computational power and practical considerations preclude efficient model analysis (fig. S11). A more granular search of *.λ* values at the selected number of components (*R* = 15) led to best fit sparsity coefficients of *.λ* = 15.0 for *Prochlorococcus* and *.λ* = 10.0 for *Synechococcus* (fig. S10, C and D), based on the maximum cross-validated FMS (see Methods). The *Prochlorococcus* model parameters (*R* = 15, *.λ* = 15.0) resulted in a cross-validated SSE of 0.84 and a cross-validated FMS of 0.54, and the *Synechococcus* model parameters (*R* = 15, *.λ* = 10.0) resulted in an SSE of 0.92 and an FMS of 0.53. The models fit with the selected parameters were then analyzed to evaluate the consistency and robustness of the resulting components.

### Evaluation of component robustness

The FMS calculated during model fitting measures component consistency in the model as a whole. However, this metric does not measure the consistency and robustness of individual components. We therefore extracted and aligned components across 300 bootstrapped models to evaluate consistency, robustness to alternate parameters, and the percentage of variation attributable to each individual component (Fig. 3). In each instance, the component-specific FMS scores were calculated based on comparison to a reference model parameterized with the identified best fit parameters (see Materials and Methods). Among the models with best fit parameters, a subset of components exhibited a tight distribution of component-specific FMS scores across bootstraps, indicating a high level of consistency between sample replicates (Fig. 3, B and E). The majority of components exhibited a median FMS of 0.5 or greater, including 13 of the 15 *Prochlorococcus* components (Fig. 3B), and 12 of the 15 *Synechococcus* components (Fig. 3E). We also aligned components from models fit with suboptimal values of *R* and compared them against the best fit model components, to assess the robustness of components to alternate *R* parameters (Fig. 3, A and D). Most components tended to exhibit a consistent median FMS score across the full range of *R* parameters in which they were detected. In general, components that accounted for a higher percentage of variation in the data (Fig. 3, C and F) tended to be the “earliest” identified, detected in models with a small *R* parameter. Components that accounted for a lower percentage of variation tended to be picked up only by the more flexible models parameterized with a greater number of components. Regardless of the minimum *R* parameter at which they were first detected, most components were subsequently detected at all values of *R* greater than this minimum, in an increasing percentage of bootstraps, and with a consistent FMS. This result indicated model stability, and suggested that the patterns represented by these components are robust to model mis-specification.

**Fig. 3.**
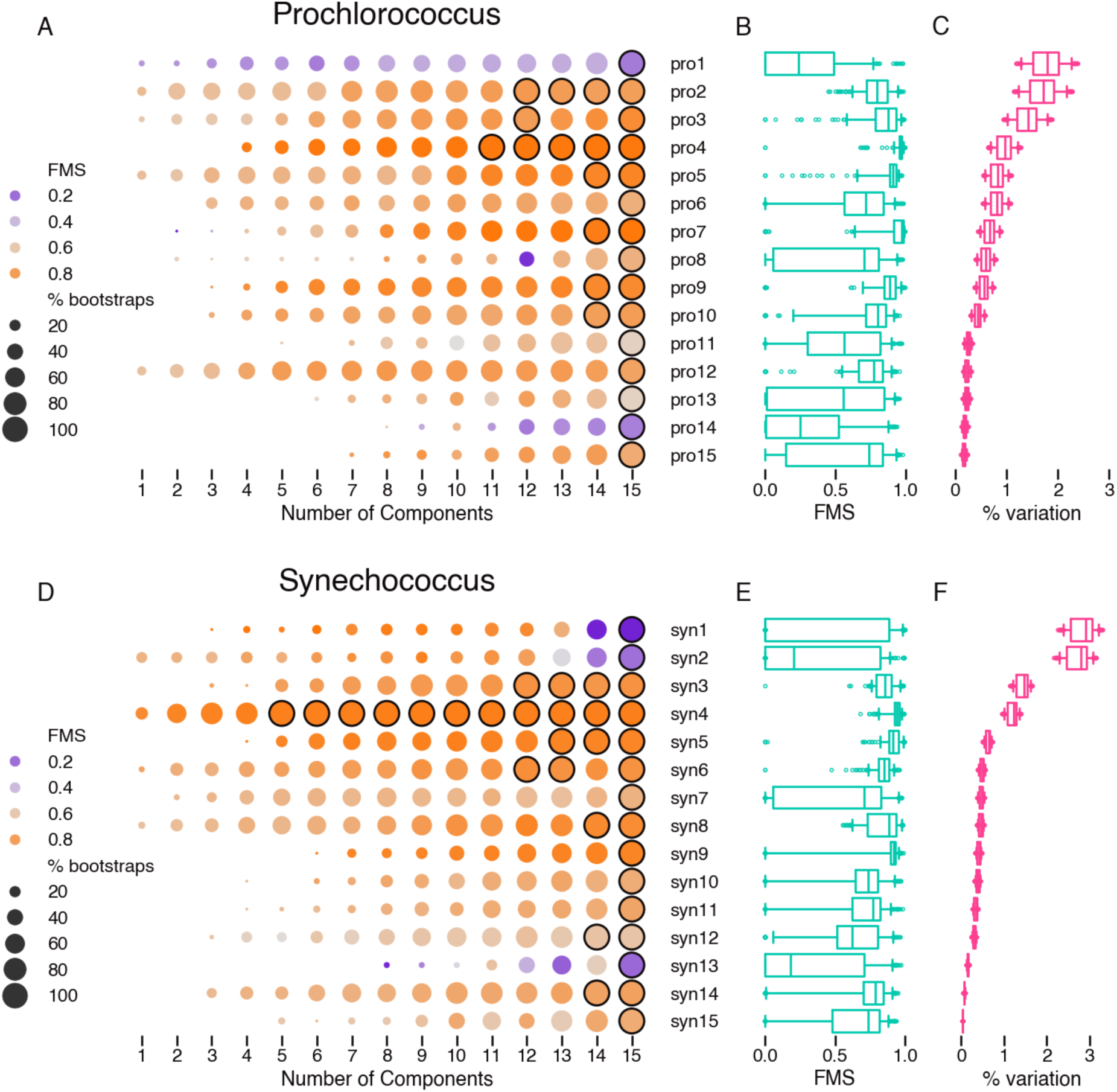
Individual components are robust to alternate values of *R* parameter. Analysis of components from sparse tensor decomposition models fit to (A-C) *Prochlorococcus* and (D-F) *Synechococcus* residual transcript abundance data. (A, D) Median factor match score (FMS) of components from models fit with different numbers of components (*R*) versus their corresponding best match component in the reference model with best fit parameters. Circle size represents the percentage of 30 bootstrap models found to contain the indicated component and circles outlined in black show components that were present in every bootstrap. (B, E) Distribution of component FMS across 300 bootstraps of best fit models. (C, F) Distribution across 300 bootstraps of percent variation represented by each component of best fit models. Boxes show inner 50th percentile centered on the median, whiskers indicate 5th and 95th percentiles, and outlying points are plotted individually.

We also evaluated the components for robustness to alternate parameterization of the spar-sity coefficient. Components exhibiting a high median FMS with best fit parameters tended to maintain consistently high FMS scores across a wide range of alternate *.λ* values (fig. S12). Although the sizes of the gene clusters naturally differ across different values of *.λ*, for most components a consistency of FMS scores across sparsity coefficients spanning almost two orders of magnitude indicated a stability in the signal modeled (fig. S13). This reinforces the validity of our sparse modeling approach and suggests that the signal of many components is dominated by a core set of co-expressed genes that can be identified by applying an optimal level of sparsity. Based on these results we chose to focus the remainder of our analysis on the 13 *Prochlorococcus* components and 12 *Synechococcus* components that fell above a median FMS threshold of 0.5, removing components pro1, pro14, syn1, syn2, and syn13 from further analysis.

### Interpretation of components

A cluster weight profile was generated for each of the remaining robust components by compiling the bootstrapped gene (CyCOG), taxon (clade), and sample weights (data S1 and S2). The profile delineates gene cluster membership, and summarizes the taxonomic and spatiotemporal patterns of cluster activity. At a 50% bootstrap threshold, the minimum *Prochlorococcus* cluster size was 4 CyCOGs, the maximum 100 CyCOGs, and the median cluster consisted of 22 unique CyCOGs. Of the 5,023 CyCOGs detected in the *Prochlorococcus* pangenome, 404 CyCOGs (*⇠* 8%) were represented in at least one cluster. In *Synechococcus*, cluster size ranged from 16 to 107 CyCOGs with a median of 65 CyCOGs, altogether representing 638 out of 6,478 CyCOGs (*⇠* 10%) detected in the *Synechococcus* pangenome. These statistics suggest that for both the *Prochlorococcus* and *Synechococcus* populations in this dataset, the most dramatic changes in expression are attributable to less than 10% of the expressed pangenome, perhaps corresponding to CyCOGs with a critical role in acclimating to specific environmental stresses encountered in our study.

The function of each CyCOG was inferred based on the majority annotation of member proteins, using several databases: the Kyoto Encyclopedia of Genes and Genomes (KEGG), Clusters of Orthologous Genes (COG), Pfam, and Tigerfam. Of the total 897 CyCOGs represented in at least one *Prochlorococcus* or *Synechococcus* cluster, we could assign a predicted function to 642 CyCOGS, while 255 CyCOGs had no assigned function. These CyCOG annotations, in conjunction with the KEGG pathway database, were used to analyze each cluster for over-represented biological functions (see Materials and Methods). All clusters were significantly enriched for at least one of 113 represented pathways, with the median cluster enriched for 12 pathways (data S3). These results signify biological coherence in the gene composition of each cluster, which we next explored in the context of the compiled taxon and sample weights.

The cluster weight profiles, combined with CyCOG annotations and sample metadata, provide an intuitive interpretation of Barnacle components, as illustrated by *Synechococcus* component syn8 (Fig. 4). Component syn8 encompassed 49 CyCOGs with nonzero median weights, and the four CyCOGs with the largest weights were detected in more than 95% of bootstraps (Fig. 4A). Component syn8 also included six CyCOGs with negative median weights, demonstrating how Barnacle can incorporate within the same cluster genes with positively and negatively correlated expression profiles. The syn8 CyCOGs included genes involved in photosynthesis, carbon fixation, and adaptation to scarce iron (data S2). The taxon weights exhibited a tight distribution across bootstraps, indicating a consistent pattern in *Synechococcus* clades I and IV, and to a lesser degree clades CRD1, CRD2, and II (Fig. 4B). The dataset incorporated samples collected under four different experimental protocols across the three cruises (fig. S1), and cluster syn8 exhibited anomalous expression in each sample group (Fig. 4, C to F). In all three cruises the elevated expression of syn8 in surface waters was confined to the northern latitudes sampled, with little anomalous expression detected south of 32°N (Fig. 4C). In 2019, a diel study at 42°N followed the same mass of water for 60 hours, sampling every 4 hours. Component syn8 exhibited a clear circadian pattern in this dataset, with a peak in expression each day at 6 am, and a nadir between 2 pm and 6 pm (Fig. 4D). In depth profiles collected in 2017 and 2019, syn8 exhibited elevated expression at the latitudes where surface expression was also detected, and weights generally decreased with increasing depth (Fig. 4E). In 2017, two sets of on-deck incubation experiments were conducted at three different latitudes. In the first set of experiments, 20 liter seawater samples were supplemented with nitrogen, phosphorous, and/or iron, and samples were collected for metatranscriptome sequencing after 96 hours of incubation. In the second set, 4 liter seawater samples were incubated with one of three sulfur-containing compounds and samples were collected for metatranscriptome sequencing after about two hours. Cluster syn8 expression was unremarkable in the 4 liter incubations (data S2), and in the experiments conducted at 33°N (Fig. 4F). However, in the incubation experiments conducted at 41°N, the syn8 cluster showed elevated expression in both the 0-hour and 96-hour control treatments, whereas no anomalous expression was detected in the three nutrient addition treatments, all of which included iron. Weights showed a wider spread across bootstraps in the transition zone experiments conducted at 37°N, indicating increased variability between these replicates. In this experiment the syn8 cluster showed elevated expression in the 0-hour and 96-hour controls, as well as the treatment amended with nitrogen and phosphorous, but no iron. In the two 96 hour treatments in which iron was added, syn8 showed no anomalous expression. In summary, the cluster profile of syn8 models a set of genes potentially related to photosynthesis, carbon fixation, and acclimation to low iron, common to *Synechococcus* clades I, IV, CRD1 and CRD2, with a joint pattern of anomalous expression in the northern part of the transect that exhibited a circadian peak at 6 am and was suppressed with the addition of iron.

**Fig. 4.**
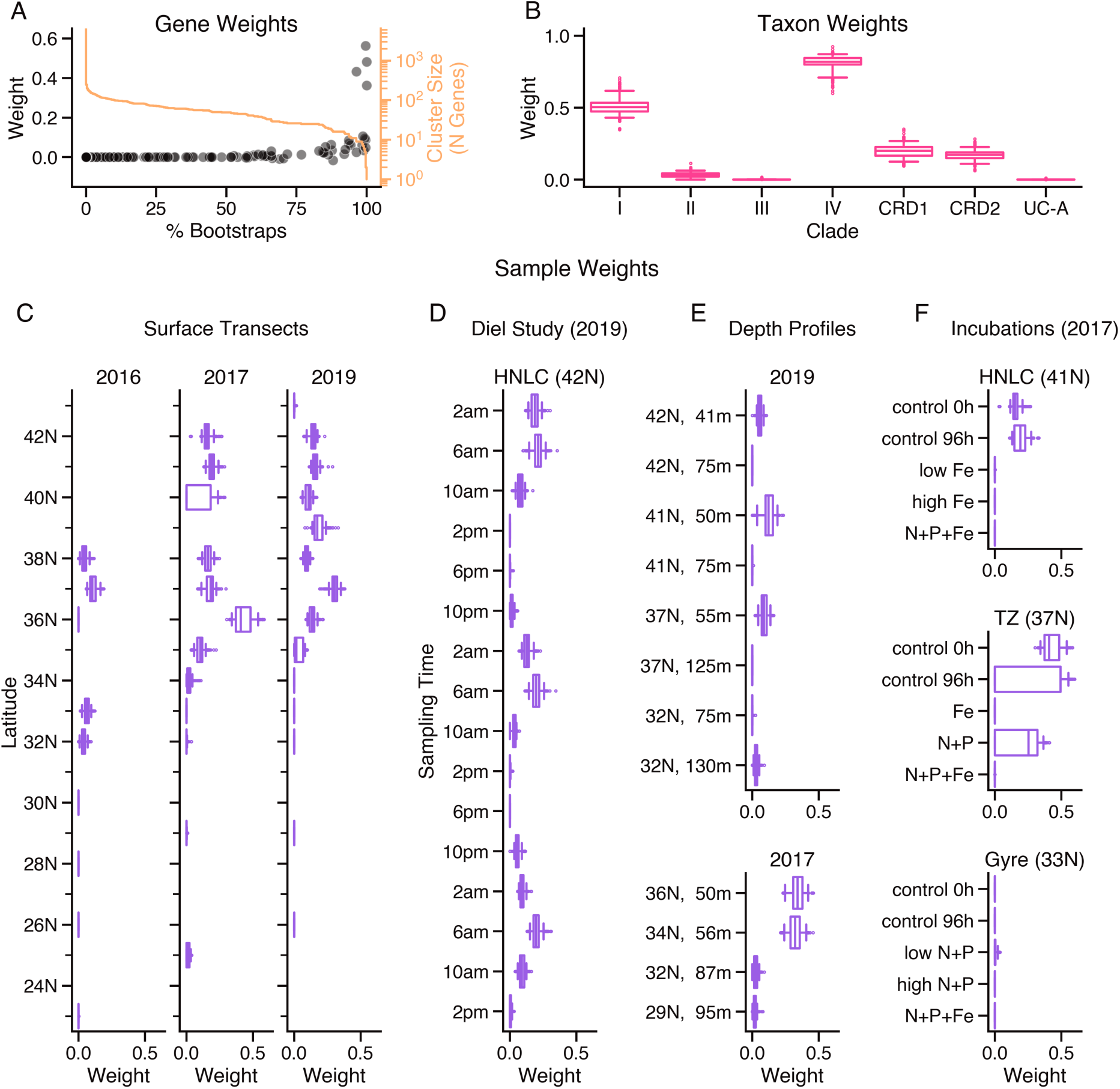
Example cluster profile of *Synechococcus* component syn8. (A) Median bootstrapped gene weight (black circles) of each CyCOG versus percentage of 300 bootstraps in which the CyCOG was included in component syn8 with a non-zero weight. Cluster size (number of CyCOGs) as a function of percent bootstrap threshold (orange line). Distribution of (B) taxon weights and (C-F) sample weights across 300 bootstraps. Boxes show inner 50th percentile centered on the median, whiskers indicate 5th and 95th percentiles, and outlying points are plotted individually. (C) Surface samples (≤15 meters) collected across 3 cruise years, binned by latitude. (D) Diel study samples binned by collection time of day. (E) Depth samples (*>*15 meters) collected in 2017 and 2019. (F) Incubation experiments in which indicated nutrients were amended to 20 liter seawater samples and transcriptomes collected after 96 hours. Component syn8 showed no significant activity in additional 4 liter incubations collected after 2 hours (data S2).

### Comparison between cluster weight profiles

To determine whether different components modeled similar processes, we compared cluster weight profiles between all pairs of *Prochlorococcus* and *Synechococcus* components (Fig. 5 and fig. S14). A majority of components corresponded to clusters of distinct, non-overlapping sets of genes (Fig. 5A). Four groups of clusters (highlighted in color) had correlated gene weight profiles. Each of these clusters was specific to different clades (Fig. 5B). For example, the gene weight profiles of clusters syn7, pro6, syn4, and syn10 represented similar sets of CyCOGs and yet their taxon weights differed: syn7 was primarily specific to *Synechococcus* clade CRD2, syn4 to clades I and IV, and syn10 to clades CRD1 and CRD2, whereas pro6 was specific to the HLI clade of *Prochlorococcus*. Several clusters had evident circadian expression patterns in the 2019 diel study samples, including syn8 (Fig. 4), syn4, and syn6 (data S2). Circadian expression was more difficult to discern for members of the other clusters, and we therefore inferred a daily peak expression time for each cluster by fitting a kernel density estimate to aggregated sampling times, weighted by the median component sample weights (fig. S15). The first group of four clusters (syn7, pro6, syn4, syn10) exhibited a peak expression time around dusk, whereas the other six clusters (pro5, syn8, pro7, syn8, pro15, and syn12) had a peak expression time around dawn (Fig. 5C).

**Fig. 5.**
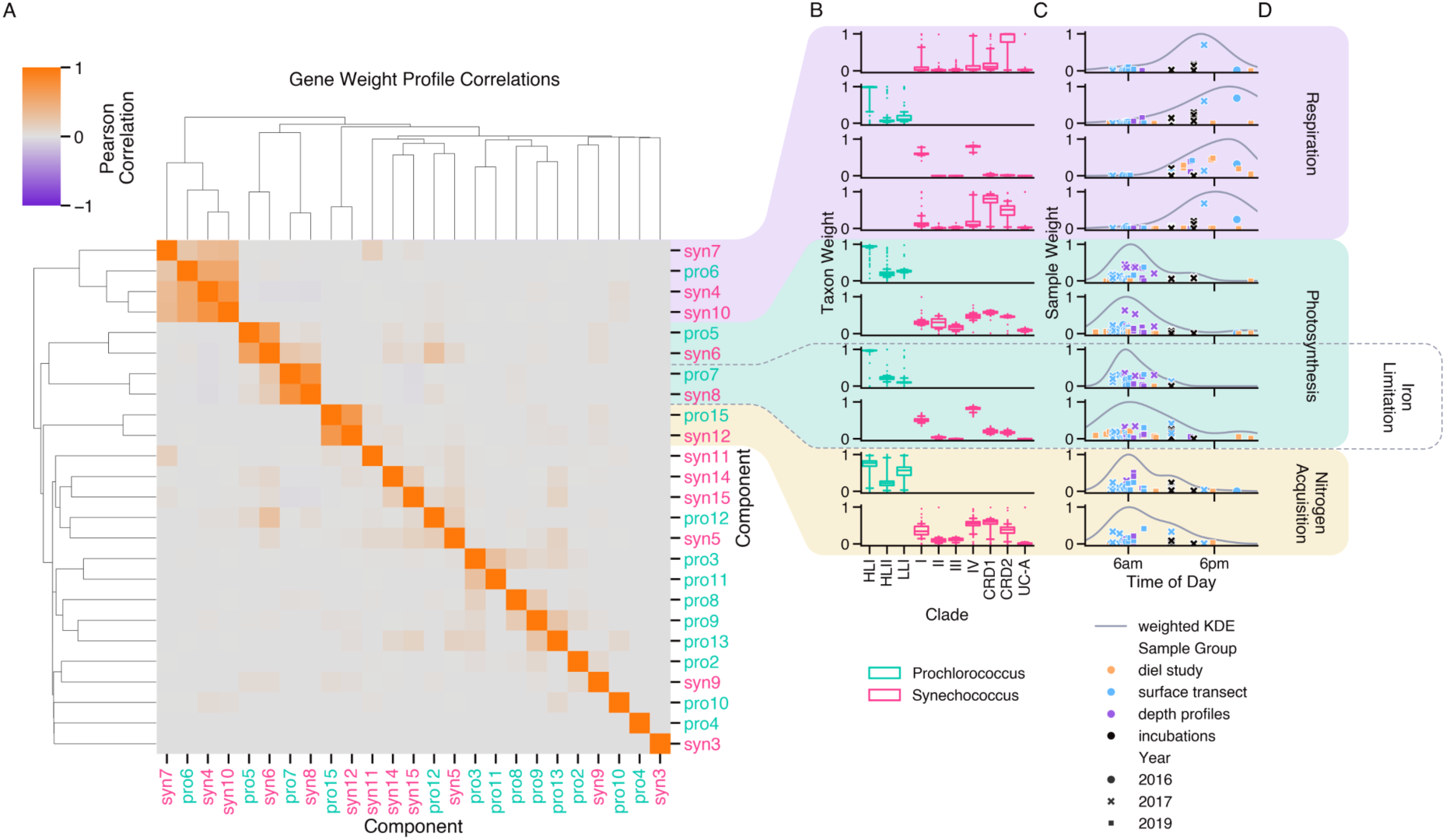
Subsets of *Prochlorococcus* and *Synechococcus* clusters exhibit similar gene content and temporal expression profiles. (A) Hierarchically clustered heatmap of pairwise Pearson correlations comparing gene weight profiles of *Prochlorococcus* and *Synechococcus* model components. Subsets of clusters with similar gene weight profiles are highlighted. (B) Distribution of bootstrapped taxon weight profiles of highlighted clusters. (C) Sample weights of highlighted profiles plotted by time of day of collection. In grey a weighted kernel density estimate indicates daily peak expression time. (D) Summary of predominant physiological processes of highlighted clusters as determined by enriched KEGG pathways and gene annotations.

Common physiological processes were apparent within each of the groups of clusters (Fig. 5D). The clusters in the first group (syn7, pro6, syn4, and syn10) were each enriched in two KEGG pathways involved in aerobic respiration: oxidative phosphorylation and the pentose phosphate pathway (data S3). These clusters also had positive gene weights for all three subunits and several assembly proteins of cytochrome c oxidase, indicating elevated expression of the primary terminal oxidase. Two groups of clusters (pro5 and syn6, pro7 and syn8) were enriched in KEGG pathways describing carbon fixation in photosynthetic organisms, the pentose phosphate pathway, and glycolysis/gluconeogenesis (data S3). The KEGG pathway describing photosynthesis was also enriched in clusters pro5 and syn6. Photosynthesis was not enriched in pro7 and syn8, however syn8 showed increased expression of two photosystem II binding proteins, PsbQ and Psb28 (data S2). In addition, high gene weights were assigned to the iron stress inducible genes *isiA* and *isiB*, both of which are understood to interact with photosynthesis processes (*25–28*). The fourth group consisted of clusters pro15 and syn12, which were both enriched for processes involved in nitrogen acquisition and assimilation, including ABC transporters in both clusters and nitrogen metabolism in syn12 (data S3). Although cluster pro15 was not enriched for nitrogen metabolism, the four CyCOGs with nonzero median weights in the cluster were annotated as components of ammonium, urea, and taurine transporters (data S1). In summary, although the majority of clusters modeled distinct cellular processes, two independent analyses reiterated similar patterns of anomalous expression in gene modules related to respiration, photosynthesis, nitrogen acquisition, and acclimation to scarce iron, indicating prominent, parallel shifts in these physiological activities across multiple co-occurring cyanobacterial populations in the North Pacific.

### Comparison to laboratory gene expression clusters

To evaluate whether the co-expression patterns detected in field data recapitulated patterns seen in laboratory studies, we next compared the Barnacle-identified *Prochlorococcus* and *Synechococcus* clusters against previously identified clusters from a diel study of *Prochlorococcus* strain MED4, grown under nutrient replete culture conditions (Zinser et al., 2009) (*29*). We mapped the MED4 genes to their corresponding CyCOG (data S4), and calculated a weighted F1 score that compared the gene content of each pair of Zinser- and Barnacle-identified clusters in an all-by-all manner. After correcting for multiple comparisons, 4 of the 18 Zinser clusters overlapped significantly with 9 of the 25 Barnacle clusters (fig. S16). The four respiration clusters pro6, syn4, syn7, and syn10 (Fig. 5) were significantly similar to Zinser cluster 5, which Zinser et al. found to be enriched for respiratory terminal oxidases and degradation of proteins, peptides, and glycopeptides. The peak expression time of the Barnacle respiration clusters occurred around 6 pm (Fig. 5C) and the Zinser cluster displayed a peak expression time of 5:30 pm *±* 0.4 hours, in both instances around sunset. Barnacle photosynthesis clusters pro5, syn6, and syn8, along with cluster pro12 displayed a peak expression time around 6 am (Fig. 5 and fig. S15) and were significantly similar to Zinser cluster 16 (fig. S16), which had a peak expression time of 5:30 am *±* 0.5 hours and was enriched for ATP synthase and CO_2_ metabolism. Barnacle clusters pro12, syn6, and syn9 significantly overlapped with Zinser cluster 13 (fig. S16), a cluster enriched for ribosomal proteins with a peak expression time of 3:24 am *±* 0.4 hours. The most significantly enriched KEGG pathway in these three Barnacle clusters corresponded to ribosomal proteins (data S3) and the peak expression time was approximately 6 am (fig. S15). The second most significantly enriched KEGG pathway in pro12 corresponded to photosynthesis (data S3). The pro12 cluster also overlapped significantly with Zinser cluster 1 (fig. S16), a cluster enriched for photosystems I and II and with a peak expression time of 8:18 am *±* 0.7 hours. The remarkable similarity in gene content and expression peaks between the Barnacle clusters identified from environmental data and the clusters identified in laboratory-grown MED4 reiterates the power of both our modeling approach and laboratory studies with model organisms.

### Acclimation to nutrient scarcity in the North Pacific

The North Pacific Ocean is characterized by iron limitation of photosynthetic organisms in subarctic regions and nitrogen limitation in the subtropical regions (*19*). Our comparison of cluster gene weight profiles revealed one pair of clusters related to nitrogen acquisition (pro15, syn12) and a second pair of clusters (pro7, syn8) containing genes linked to iron stress (Fig. 5). Within each pair, the two clusters shared similar sample weight profiles (fig. S14), with pro7 and syn8 showing larger weights in the northern section of the transect (Fig. 4 and data S1) and pro15 and syn12 showing larger weights in the southern section (data S1 and S2). These observations led us to deduce that the four clusters model processes relevant to acclimation and ecological partitioning along nutrient gradients in the North Pacific, providing an opportunity to study the in situ physiological effects of these environmental stresses by examining cluster gene content and sample weights in more detail (Fig. 6).

**Fig. 6.**
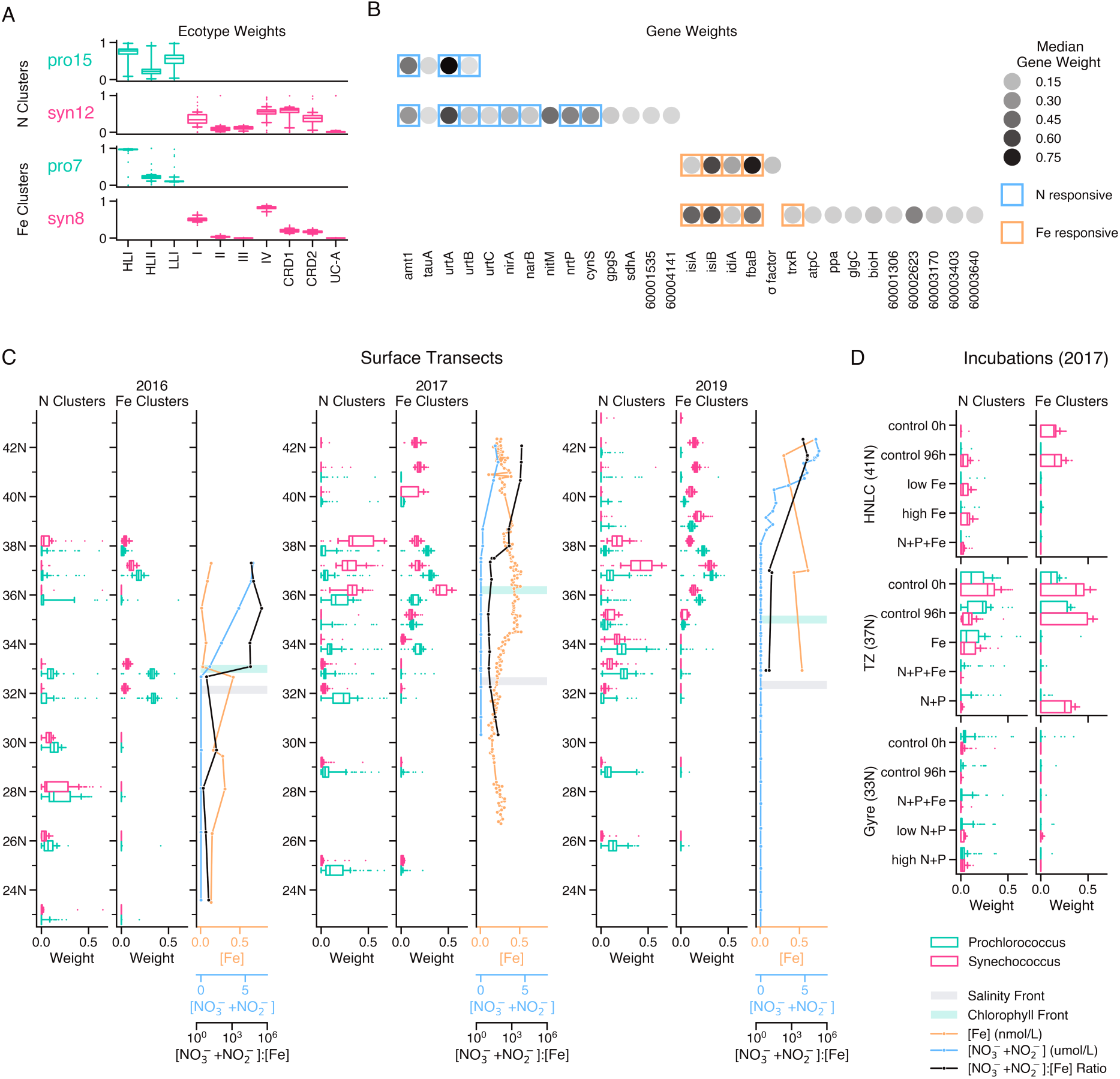
Independent *Prochlorococcus* and *Synechococcus* clusters signal acclimation to scarce nitrogen and iron overlapping in the subtropical-subarctic transition zone. Bootstrapped component weight profiles of *Prochlorococcus* (green) and *Synechococcus* (pink) clusters enriched for genes related to nitrogen scarcity (pro15, syn12) and iron scarcity (pro7, syn8). (A) Distributions of cluster taxon weights. (B) Top cluster genes with absolute median weight *2::* 0.04. Boxes indicate genes previously reported to have significantly altered expression in response to nitrogen deprivation (blue) (*32, 33, 35*) and iron deprivation (orange) (*25, 40–43*). (C) Surface sample weight distributions for three years of cruises, binned by latitude, with nitrogen-responsive cluster weights in leftmost columns and iron-responsive cluster weights in middle columns. Rightmost columns show the nitrogen-to-iron ratio (black) calculated using measurements of surface dissolved iron concentration (orange) and dissolved nitrate plus nitrite (blue). Salinity and chlorophyll fronts as reported by Juranek et al. (2020) are indicated by grey and green bars, respectively (*80*). (D) Distributions of sample weights from 20 liter nutrient amendment experiments measured after 96 hours of incubation and conducted at three latitudes in 2017. Boxes show inner 50th percentile centered on the median, whiskers delineate 5th and 95th percentiles, and outlying points are plotted individually.

#### Nitrogen acclimation response

*Prochlorococcus* and *Synechococcus* populations in the North Pacific uptake ammonium and urea as their primary nitrogen source (*30*), and subpopulations of both genera have been shown to assimilate nitrate (*30, 31*). A close examination of the CyCOGs in clusters pro15 and syn12, and their associated gene weights (all of which were positive), revealed elevated expression of four genes common to both clusters: *amt1*, *tauA*, *urtA*, and *urtB* (Fig. 6B). The *urtA* and *urtB* genes encode two components of a urea ABC-transporter system and *amt1* encodes an ammonium transporter. Increased expression of *amt1* and *urtABCDE* has been observed in response to nitrogen deprivation in marine *Prochlorococcus* (*32*) and *Synechococcus* (*33*). The *tauA* gene encodes the substrate-binding component of an ABC-transporter system that has recently been proposed to import guanidine in marine cyanobacteria from low nitrogen environments (*34*). The *Prochlorococcus* cluster pro15 was limited to these four CyCOGs, whereas the *Synechococcus* cluster syn12 consisted of a total of 20 CyCOGs (data S5). The additional syn12 CyCOGs included nitrite reductase (*nirA*), nitrate reductase (*narB*), the formate/nitrite transporter (*nitM*) and the bi-specific nitrate/nitrite permease (*nrtP*), all of which are involved in nitrate and nitrite assimilation (Fig. 6B). In *Synechococcus* the expression of *nirA*, *narB*, and *nrtP* is controlled by the universal nitrogen response regulator NtcA, and up-regulated in response to nitrogen deprivation (*35*). Cyanate lyase (*cynS*), included in syn12, is also expressed in response to ammonium deprivation in an NtcA-dependent manner (*36*). A key enzyme in the synthesis of glucosylglycerate, *gpgS*, was also included in syn12. Glucosylglycerate is a compatible solute that has been found to accumulate in *Synechococcus* under nitrogen limitation, possibly replacing glutamate as a negatively charged counterion (*37*). We detected additional genes related to nitrogen metabolism with bootstrap values that fell below the 50% threshold, including CyCOGs corresponding to NtcA (CyCOG 60000127) and glutamine synthase (CyCOG 60000563) (data S1 and S2). A small fraction of pro15 bootstraps also included nitrate and nitrite reductases (data S1), possibly reflective of the patchy distribution of *nirA* and *narB* among *Prochlorococcus* strains at the sub-clade level (*31*).

#### Iron acclimation response

A deeper evaluation of the gene content of clusters pro7 and syn8 revealed four shared CyCOGs with large positive weights, all of which are implicated in acclimation to iron scarcity in cyanobacteria (Fig. 6B). Three of the shared CyCOGs are canonical markers of iron stress: *isiA* encodes a chlorophyll binding protein that forms an antenna structure around photosystem I in iron-limited cyanobacteria (*25, 27, 28*), *isiB* encodes flavodoxin, an iron-free analog of the iron-containing electron transfer protein ferredoxin (*25, 26*), and *idiA* is hypothesized to encode either a protein that protects photosystem II against oxidative stress during iron limitation (*38*) or a component of an ABC-type Fe(III) transporter (*39*). All three genes have been shown to be up-regulated in response to iron limitation in both *Prochlorococcus* (*40, 41*) and *Synechococcus* (*25, 42, 43*). The fourth shared gene, *fbaB*, encodes a class I fructose bisphosphate aldolase (FBA), an enzyme involved in several carbon metabolism pathways including carbon fixation. There are two variants of FBA: class II FBAs require a divalent iron or zinc cation as a cofactor, whereas class I FBAs do not require a metal cofactor (*44*) and have been found to be up-regulated in response to iron limitation in *Synechococcus* (*43*) and diatoms (*45*). The class I FBA (*fbaB*; CyCOG 60001290) was assigned positive weights in both pro7 and syn8, indicating elevated expression, whereas the class II FBA (CyCOG 60001287) was not included in either cluster.

As with the nitrogen-related clusters, the iron-related *Synechococcus* cluster (syn8) was larger than the iron-related *Prochlorococcus* cluster, encompassing a total of 49 CyCOGs, of which 17 are uncharacterized (data S5). The annotated CyCOGs specific to *Synechococcus* that showed the highest degree of elevated expression (Fig. 6B) fell into one of four functional categories: oxidative stress response (*trxR*), ATP production (*atpC*, *ppa*), glycogen production (*glgC*), and biotin biosynthesis (*bioH*). The gene product of *trxR* is thioredoxin reductase, an enzyme involved in mitigating oxidative stress that has shown increased expression in *Synechococcus* subjected to low iron conditions (*43*). To a lesser degree, the antioxident peroxiredoxin (*ahpC*) also showed elevated transcription in syn8 (data S5); elevated levels of peroxiredoxin have been observed in the proteome of iron-limited cultures of clade II *Synechococcus* (*46*). Similarly, in addition to *glgC*, the genes *glgA* and *glgB* were included in syn8 with small positive weights, meaning the complete glycogen synthesis pathway exhibited increased expression in syn8. Increased glycogen production has been observed in laboratory studies of *Synechococcus* facing iron scarcity (*47*), and may be connected to stress response in cyanobacteria (*48*). Finally, biotin biosynthesis requires the precursor pimelate thioester, which can be synthesized through either an iron-dependent (*bioI-bioW*) or an iron-independent (*bioC-bioH*) pathway (*49*). The elevated expression of *bioH* in cluster syn8 is consistent with a pattern of swapping out iron containing proteins with iron-free analogs during iron limitation, and may reflect another instance of this strategy to reduce the cellular iron quota of *Synechococcus*.

Both the pro7 and syn8 clusters included additional CyCOGs that exhibited a smaller degree of differential expression and yet were consistently represented in bootstraps (data S5), thus providing additional support for trends inferred from the CyCOGs with larger gene weights. Two additional CyCOGs potentially involved in iron acquisition had positive gene weights: CyCOG 60002762 in syn8 was annotated as a hemoglobin/transferrin/lactoferrin receptor protein (KEGG Orthology K16087), and *gap2* in pro7 encodes glyceraldehyde-3-phosphate dehydrogenase (GAPDH). Although GAPDH is primarily involved in the Calvin cycle, it has also been shown to moonlight as a siderophore receptor in diverse bacteria (*50*). In addition to *gap2*, a number of other Calvin cycle genes showed elevated expression in both clusters: *rpe* in pro7, and in syn8, *rbcS*, *csoSCA*, *csoS2*, *prkB*, and *glpX*. In syn8 the increased expression of carbon fixation genes was accompanied by elevated expression of photosystem II binding proteins (PsbQ, Psb28) and genes involved in phycobilin biosynthesis (*hemE*, *hemH*, *hmuO*), indicating that the *Synechococcus* iron scarcity response includes an adjustment of several structures and pathways related to photosynthesis. The syn8 cluster also included genes that displayed negative gene weights: three cytochrome c oxidase genes, indicating decreased expression of the respiratory terminal oxidase, and the chlorophyll biosynthesis gene *clhN*. Cyanobacteria encode two protochlorophyllide reductases; *clhN* encodes the iron-sulfur subunit of one while its analog has no iron requirement (*51*). Finally, in line with an elevated oxidative stress response, syn8 included increased expression of an under-characterized CyCOG (60003403) that shares homology with DoxX, a component of a membrane-bound protein complex that co-localizes a thiol oxidoreductase with a superoxide dismutase in *M. tuberculosis* (*52*). Altogether the gene weights of clusters pro7 and syn8 suggest a broad program of transcriptional modulation to import more iron, decrease cellular iron quota, adjust photosynthesis, and mitigate oxidative stress.

#### Latitudinal trends in nutrient acclimation

Surface sample weights indicated that the transcriptional patterns modeled by clusters pro7, syn8, pro15, and syn12 were geographically bounded, consistent across study years, and echoed in both *Prochlorococcus* and *Synechococcus* populations (Fig. 6C). The nitrogen acquisition clusters pro15 and syn12 showed consistent low levels of elevated expression in samples collected in the subtropical gyre and further north, to about 38°N (Fig. 6C). In contrast, the iron acclimation clusters pro7 and syn8 exhibited elevated expression north of the subtropical gyre, with syn8 expression extending to around 42°N, the northernmost stations sampled on the 2017 and 2019 cruises (Fig. 6C). The *Prochlorococcus* iron acclimation cluster pro7 was not detected in the northernmost surface samples, likely because this cluster primarily represents the HLI clade (Fig. 6A), which did not make up a significant portion of the *Prochlorococcus* community in those samples (fig. S6). Rather, at these latitudes the *Prochlorococcus* community was dominated by the LLI and LLVII clades, the former noteworthy for a high frequency of genes involved in iron acquisition via siderophore uptake (*53*). In the transition zone (TZ) between the subtropical and subarctic gyres, expression of both the nitrogen and iron acclimation clusters was elevated in both *Prochlorococcus* and *Synechococcus* populations (Fig. 6C). The ratio of dissolved nitrate to dissolved iron increased north of 33°N in 2016 and north of 37°N in 2017 and 2019. In all three years this inflection point coincided with the region in which the iron and nitrogen acclimation clusters showed simultaneous elevated expression in both the *Prochlorococcus* and *Synechococcus* populations.

To investigate whether the in situ gene expression patterns observed were consistent with experimental evidence of nutrient limitation, we examined sample weights associated with the 2017 nutrient amendment incubation experiments (Fig. 6D). At the gyre station (33°N), the nitrogen clusters (pro15, syn12) had non-zero sample weights, indicating a low level of elevated expression, whereas the weights for the iron acquisition clusters (pro7, syn8) were zero. Conversely, at 41°N the *Synechococcus* iron cluster syn8 showed elevated expression in both the 0-hour and 96-hour controls, whereas the *Synechococcus* nitrogen cluster syn12 was associated with a zero weight in the 0-hour control sample and slightly elevated expression in the 96hour incubations, although less so in the treatment amended with supplemental nitrogen. The *Prochlorococcus* clusters showed no activity in this set of samples since *Prochlorococcus* clade HLI was not abundant at this latitude (fig. S6). In the transition zone incubations conducted at 37°N, all four clusters showed elevated expression in the 0-hour and 96-hour controls (Fig. 6D). Moreover, at this site treatments amended with nitrogen resulted in sample weights of zero for the nitrogen clusters, and treatments amended with iron resulted in sample weights of zero for the iron clusters. These results imply that expression of clusters pro7 and syn8 was specifically suppressed with the addition of iron, while pro15 and syn12 expression was suppressed with the addition of nitrogen. Furthermore the results provide evidence of a North Pacific gradient of nitrogen limitation in the subtropical gyre and iron limitation in the subarctic gyre, and indicate a region of simultaneous nutrient stresses experienced by the *Prochlorococcus* and *Synechococcus* populations at the transition zone boundary between these two ecosystems.

## Discussion

Although metatranscriptome sequencing is now common in studies of environmental microbiomes, the physiological states reflected in a metatranscriptome often remain obscured by data complexity that exceeds the capacity of traditional analytical techniques and limited prior knowledge of functional gene networks. Barnacle is an unsupervised signal discovery tool, tailored to metatranscriptomic datasets, that we developed to help bridge this technological gap and to investigate patterns of gene expression that determine the ecology of a microbiome. The core of the method, tensor decomposition, reflects the inherent multiway structure of metatranscriptomic datasets. It also serves as a fundamental constraint on the gene clustering problem (*54*), enabling resolution of overlapping expression patterns, even when obscured by high noise (Figs. 1 and 2). We prioritized robust, interpretable inferences by imposing nonnegativity and sparsity constraints, and used sample replicates to implement cross validation and bootstrapping, enabling us to quantify confidence in the association of each gene with each component. By combining this methodology with the variance stabilizing transform for normalization (*23, 24*), we focused our analyses towards population-level shifts in transcription. The normalization approach also addressed overdispersion in our data (fig. S8) – a common challenge in metatranscriptomic datasets (*14*) – and enabled us to distinguish between zero values caused by low sequencing depth, the absence of a gene ortholog in a taxon, or potential downregulation. Applying the methodology to cyanobacteria populations that form the foundation of North Pacific ecosystems, we demonstrated the power of our approach to uncover processes that drive in situ microbiome dynamics, including the circadian choreography of essential metabolic processes and mechanisms of acclimation to shifting nutrient concentrations.

Two of the most robust components identified in our analysis, pro7 and syn8, suggested that the acclimation response to scarce iron exhibits a similar pattern in *Prochlorococcus* and *Synechococcus*, with important physiological differences distinguishing the two lineages. As an efficient electron transfer agent, iron is essential to the functioning of many core cellular processes, including photosynthesis and respiration. Consequently, iron scarcity poses a serious obstacle to photosynthetic growth and is one of the main limiting nutrients across much of the surface ocean (*19*). Overcoming iron scarcity requires coordinated adjustment of intertwined cellular processes. Three physiological categories of cyanobacterial acclimation to iron limitation were proposed by Straus (1994) (*55*): 1) acquisition of more iron, 2) compensation via the replacement of iron-requiring proteins with iron-free analogs, and 3) retrenchment, which entails a reduction of cell structures, particularly photosystem I and other iron-rich components of photosynthetic electron transport (PET) (*56*). Additionally, iron limitation poses an increased risk of oxidative damage, in part because retrenchment and compensation decrease the number of iron centers in PET, which makes saturation of the PET chain more likely and increases the likelihood that electrons will escape and form reactive oxygen species (*57*). As such, a fourth category of cyanobacterial acclimation to low iron was later added by Michel and Pistorius (2004) to distinguish the role that some iron deficiency induced genes play in mitigating oxidative stress (*38*). All four categories of cyanobacterial response to scarce iron are reflected in the physiological states modeled by clusters pro7 and syn8. As such, our results recapitulate well-established patterns of iron stress response, and additionally reveal new potential response mechanisms and contrasts in iron physiology between *Prochlorococcus* and *Synechococcus*.

*Prochlorococcus* cluster pro7 encompassed seven CyCOGs with elevated expression (Fig. 6B, data S5), each with a putative role in iron acquisition, compensation, retrenchment, or protection from oxidative stress. Three CyCOGs correspond to well-established gene markers of iron deficiency: *isiA*, *isiB* (flavodoxin), and *idiA*. The *isiA* and *idiA* gene products are hypothesized to protect photosystems I and II from oxidative damage, respectively (*38*). The *idiA* gene product may also be involved in iron acquisition as it displays homology to the periplasmic component of the bacterial ferric uptake transporter (*38*). Flavodoxin expression is a canonical example of compensation, functionally replacing the iron-sulfur electron shuttle ferrodoxin with an iron-free analog (*40–42, 55*). The elevated expression of an iron-free fructose bisphosphate aldolase (FBA) analog in pro7 may be another example of this compensation pattern. FBAs are involved in several carbon metabolism pathways, including the Calvin cycle, as are two of the remaining CyCOGs in pro7: *rpe* encoding ribulose-5-phosphate 3-epimerase (RPE) and *gap2* encoding a glyceraldehyde-3-phosphate dehydrogenase (GAPDH). Our analysis accordingly found that carbon fixation was the most significantly enriched KEGG pathway in pro7 (data S3). The rate of carbon fixation limits the rate of photosynthesis in cyanobacteria (*58*) and so increasing the expression of critical Calvin cycle enzymes could serve to compensate for lowiron-induced retrenchment by maximizing photosynthetic efficiency. Maximizing the carbon fixation rate could also serve to reduce the formation of reactive oxygen species by maximizing the rate at which electrons flow out of PET and into the Calvin cycle, thus relieving saturation of the PET chain. Additionally, RPE employs an iron atom in its active site, and both it and GAPDH are inactivated by oxidation (*59,60*). These enzymes may be points of the Calvin cycle vulnerable to oxidative damage, and thus their up-regulation may serve as a protective response to the increased oxidative stress that accompanies iron limitation. Finally, pro7 includes one CyCOG corresponding to a sigma factor. The presence of a single sigma factor within this cluster suggests it may be involved in coordinating the *Prochlorococcus* transcriptional program across these four categories of acclimation to low iron.

*Synechococcus* cluster syn8 exhibits many of the same hallmarks of low iron acclimation seen in the *Prochlorococcus* cluster. Iron deficiency marker genes *isiA*, *isiB*, and *idiA* all show a similar degree of over-expression, as does *fbaB*, encoding the iron-free class I FBA enzyme (Fig. 6B). However, our results indicate that *Synechococcus* acclimation to low iron is more involved than that of *Prochlorococcus*: syn8 encompasses a total of 49 CyCOGs as compared to the 7 CyCOGs of pro7 (data S5). This difference may reflect the evolutionary streamlining of *Prochlorococcus* metabolism in comparison to the greater metabolic flexibility of *Synechococcus* (*61*), which allows it to thrive across a greater range of environmental conditions (*2*). *Prochlorococcus* metabolism is likely already well-equipped to cope with nutrient scarcity, and so adapting to low iron conditions requires relatively minor tuning of its transcriptional program. In contrast, while *Synechococcus* possesses the genetic potential to thrive across a range of nutrient concentrations, actualizing acclimation to low iron entails broader metabolic restructuring. Here the inferential power of Barnacle components is on full display, allowing us to detect the cohort of genes up-regulated in concert with markers of low iron stress, and to extrapolate from their collective activity the mechanism physiological acclimation in *Synechococcus*.

The shifts in gene expression specific to cluster syn8 appear primarily geared towards maximizing the efficiency of photosynthesis and downstream metabolic processes, and mitigating oxidative stress. Two CyCOGS encoding enzymes involved in oxidative stress response showed elevated expression: thioredoxin reductase (*trxR*; Fig. 6B) and peroxiredoxin (*ahpC*; data S5), both of which have been previously observed up-regulated in iron-limited *Synechococcus* (*43, 46*). Seven up-regulated CyCOGs are involved in carbon fixation, and another two are involved in ATP synthesis (data S5). Similar to RPE in *Prochlorococcus*, many of these enzymes are susceptible to oxidative damage, and their increased expression may serve as a “spare parts” repository, speeding up the repair of damaged components and reducing potential metabolic bottlenecks resulting from slow repair. Similarly, genes encoding the photosystem binding proteins PsbQ and Psb28, up-regulated in cluster syn8, are associated with photosystem II repair in cyanobacteria, following oxidative inactivation (*62, 63*). Elevated Psb28 abundance has also been observed in the proteome of clade II *Synechococcus* subjected to iron limitation (*46*). Increased expression of genes encoding the enzymes uroporphyrinogen decarboxylase (*hemE*), ferrochelatase (*hemH*), and heme oxygenase (*hmuO*) suggests an increased invest-ment in the construction of phycobilisomes, as each enzyme catalyzes a key branch-point reaction in phycobilin biosynthesis. In addition to harvesting light for photosynthesis, phycobilisomes mediate non-photochemical quenching in *Synechococcus*, dissipating excess light energy via state transitions (*64*) and interaction with the orange carotenoid protein (*65*). The investment in phycobilisome synthesis may enhance a cell’s capacity to divert light energy from PET and reduce the potential formation of reactive oxygen species. At the opposite end of electron transport, syn8 gene weights indicated increased expression of genes required for the glycogen biosynthesis pathway (*glgABC*), and decreased expression of the respiratory terminal oxidase (data S5). Glycogen metabolism is implicated in boosting the efficiency of photosynthesis and carbon fixation in *Synechococcus*, particularly during the transition from dark to light (*48*). Glycogen is also a terminal sink for photosynthetic electrons, and increasing its production may help relieve pressure on the PET chain. The decrease in respiration is best interpreted in the context of the diel pattern of the syn8 expression profile, which peaked around dawn (Fig. 4). Our data (Fig. 5) and previous studies (*29*) indicate that cyanobacterial respiration peaks at dusk; the observed down-regulation of respiration genes in syn8 may indicate a deepening of the diel segregation of photosynthesis and respiration in response to scarce iron, perhaps to minimize additional electron inputs to the components of the electron transport chain that are shared by photosynthesis and respiration (*66*). Altogether the syn8 cluster touches multiple stages of photosynthesis, from light harvesting to ATP synthesis, carbon fixation, and glycogen production, suggesting a program of acclimation to low iron in *Synechococcus* that maximizes photosynthetic electron flux, enhances diversion routes for excess reducing power, and mitigates oxidative stress. Intriguingly, the transcriptional responses in *Synechococcus* uncovered here are reminiscent of streamlining events proposed to underlie the evolutionary adaptation of *Prochlorococcus* lineages to low nutrient conditions (*61*).

At the ecosystem scale, the sample weights associated with iron acclimation clusters pro7 and syn8 showed a latitudinal trend of elevated expression in the north and lower expression in the south, inverse to the sample weights of nitrogen assimilation clusters pro15 and syn12 (Fig. 6, C and D). Just as pro7 and syn8 genes *isiA* and *isiB* have been used as indicators of low iron stress, the genes *urtABC* (urea transporter), *nirA* (nitrite reductase), and *narB* (nitrate reductase) have previously been used as indicators of low nitrogen stress (*67, 68*). Each of these genes showed elevated expression in pro15 and syn12 (Fig. 6B and data S5). Together the sample weights for these four clusters illustrate how *Prochlorococcus* and *Synechococcus* experience a progression from nitrogen scarcity in the subtropical gyre to iron scarcity in the subarctic gyre, consistent with estimates of nutrient limitation in the North Pacific surface ocean (*19*). Moreover, the cluster profiles indicate a region of overlap in the subtropical-subarctic transition zone, where the sample weights for both the iron and nitrogen acclimation clusters showed elevated expression in both *Prochlorococcus* and *Synechococcus* populations (Fig. 6C). Furthermore, in incubation studies of seawater collected from the transition zone, expression of the iron acclimation clusters was suppressed by supplementation with iron, and expression of the nitrogen acclimation clusters was suppressed by supplementation with nitrogen (Fig. 6D). This simultaneous acclimation is distinctive to the transition zone; populations in the subtropical gyre were exclusively up-regulating the nitrogen acclimation clusters whereas populations in the subarctic gyre were solely up-regulating the iron acclimation clusters. Our data are consistent with previous reports of co-limitation at ecosystem boundaries (*67, 69*), and suggest that the transition zone may be a distinct ecosystem, shaped by multiple intersecting nutrient stresses, and inhabited by distinct populations equipped to cope with these conditions.

The analytical approach developed here and applied to *Prochlorococcus* and *Synechococcus* metatranscriptomes uncovered mechanisms of acclimation that range from the function of individual genes to coordinated metabolic reprogramming to ecosystem scale processes governing the North Pacific microbiome. The unsupervised nature of our method alleviated dependency on functional annotations, enabling the concordant analysis of novel, uncharacterized gene families. This capability is critical as new surveys of environmental gene sequences routinely uncover a majority with no resemblance to previously described gene families (*8*). In this study, we found that 255 of the 897 CyCOGs included in clusters were uncharacterized, and in *Prochlorococcus* cluster pro10, the function of 10 of the top 11 CyCOGs is unknown (data S1). However, Barnacle offers a window of insight into the roles of these unknowns via their association with co-expressed, better characterized genes. In the future, this co-expression information could be combined with data on gene co-evolution, structural modeling, and genome topology to bolster computationally derived functional inferences. Finally, our approach is flexible, and could be applied to any multiway dataset that can be represented as a three-way tensor. The sparsity and non-negativity constraints make Barnacle especially well-suited for other omics datasets, including metagenomics, proteomics, and metabolomics. In these ways, the development of Barnacle, its fruitful application to North Pacific metatranscriptomes, and the resulting insights into cyanobacterial nutrient acclimation, together present a demonstration of the type of unsupervised analytical techniques that will better equip microbiome researchers to extract discoveries from complex datasets, and to discern the patterns by which molecules, cells, and ecosystems interact to drive global biogeochemical cycles.

## Materials and Methods

### Model

The sparse tensor decomposition model we developed is a constrained version of the classical CANDECOMP/PARAFAC (CP) tensor decomposition (*15, 54*). We describe the model in the context of its intended application to environmental transcript abundance data. The notation and symbols used (Table 1) adhere to those laid out by Kolda and Bader (2009) (*15*). Model implementation is available as part of the Python package Barnacle.

We consider a third-order tensor *Y* of transcript counts indexed in three modes: gene *⇥* taxon *⇥* sample. Each entry *y_ijk_* encodes the abundance of gene *i* transcripts attributed to taxon *j* in sample *k*. The length of the gene mode, *I*, reflects the number of gene orthologs measured in the dataset; the length of the taxon mode, *J*, the number of taxa; and the length of the sample mode, *K*, the number of samples. The CP model represents the data tensor *Y* as a collection of *R* components (fig. S2A), each of which models a distinct linear signal pattern present in the data, obscured by a Gaussian noise tensor *“ ⇠ N* (0, *a*^2^). That is

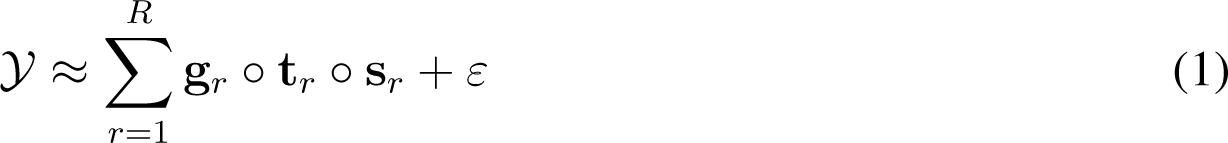

or element-wise

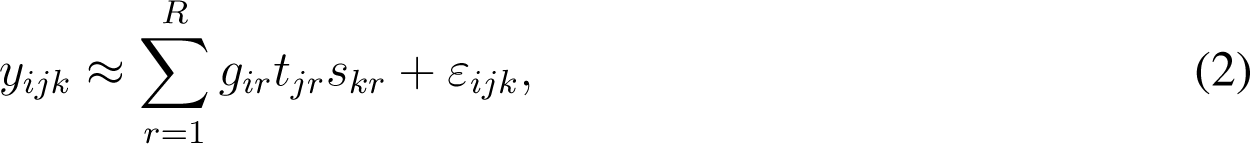

where **G** *2* R*^I⇥R^*, **T** *2* R*^J⇥R^*, and **S** *2* R*^K⇥R^* are the component matrices with columns representing the weight vectors **g***_r_*, **t***_r_*, and **s***_r_* for the gene, taxon and sample modes, respectively. Entry *g_ir_*indicates the relative contribution of gene *i* to the component *r* signal. When weight vector **g***_r_* is sparse, non-zero entries can be considered as a cluster of genes, unified by correlated expression profiles (fig. S2C). Weight vectors **t***_r_* and **s***_r_* indicate the relative activity of the component *r* signal in the modeled taxa and samples. Taking the outer product of weight vectors **g***_r_*, **t***_r_*, and **s***_r_* produces a rank-1 tensor representing component *r* (fig. S1B), and the sum of all *R* component outer products constitutes the rank-*R* tensor model, *Y*^^^.

#### Constraints

Two sets of constraints are imposed on the component matrices of the tensor decomposition model to facilitate their interpretation. First, non-negativity constraints are imposed on the taxon and sample component matrices **T** and **S**. No non-negativity constraint is imposed on the gene component matrix **G**. This combination of non-negativity constraints abrogates issues of model indeterminacy resulting from sign flips while retaining the ability to model patterns of elevated and diminished expression (positively and negatively correlated transcript abundance profiles) within the same component. Second, we promote sparsity in the gene-mode component weight vectors via the incorporation of an l1 regularization penalty applied column-wise to the gene component matrix **G**. The degree of regularization is tuned by a sparsity coefficient parameter *.λ*. This parameter indirectly controls the size of model clusters, in that a larger *.λ* will result in more zero values in the gene component matrix **G**, and thus, on average, fewer genes corresponding to non-zero values in each component. The scaling indeterminacy of CP models (component matrices in one mode can be arbitrarily scaled given an inverse scaling of the component matrices in the other modes) means that if norm-based regularization is imposed on at least one mode, then the component matrices of all modes must be regularized (*70*). As such, the gene-mode sparsity penalty is accompanied by l2 unit norm constraints applied column-wise to the taxon and sample component matrices **T** and **S**. Collectively, these constraints enable interpretation of components as clusters of genes with correlated transcript abundance patterns shared across the the taxa and samples indicated by component weights.

#### Optimization

The model is fit to data by minimizing a cost function (Eq. 3), parameterized by two userdefined variables: *R*, the number of components, and *.λ*, the sparsity coefficient of the l1 regularization penalty applied to the gene-mode. The optimal solution to the cost function is obtained by component matrices **G**, **T**, and **S** that minimize the sum of square differences between the data tensor *Y* and the model tensor (the sum of *R* components), while also minimizing the sparsity penalty on the gene component matrix. Assuming the matrices fulfill Kruskal-rank conditions (*71*), this solution is guaranteed to be unique (*18, 54*).

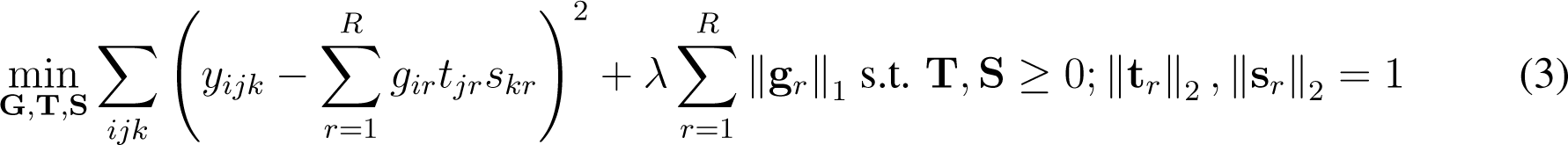

We solve the cost function using a modified version of the alternating least squares (ALS) algorithm for CP tensor decomposition (see e.g. (*15*)). Briefly, the algorithm iterates over a series of update steps, each of which cycles between the modes of the tensor, updating the component matrix of one mode while the component matrices of the other two modes remain temporarily frozen. The component matrix of each mode is updated by solving a regularized least squares regression problem, calculated using the input data tensor (unfolded along the appropriate mode) along with the Khatri-Rao product of the two frozen component matrices (Eqs. 4 to 6). The updated component matrix entries are then frozen, and the algorithm proceeds to update the component matrix of the next mode. The algorithm iteratively updates each component matrix in this manner until the change in loss between successive iterations drops below a tolerance threshold.

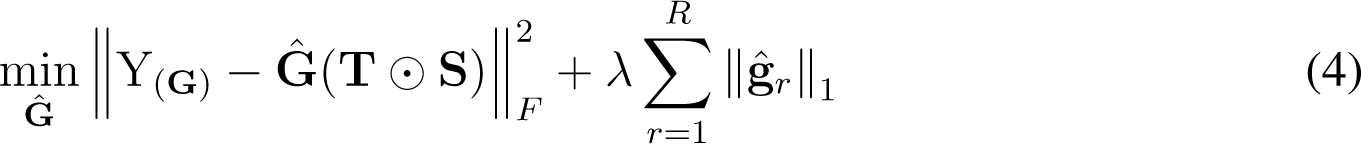

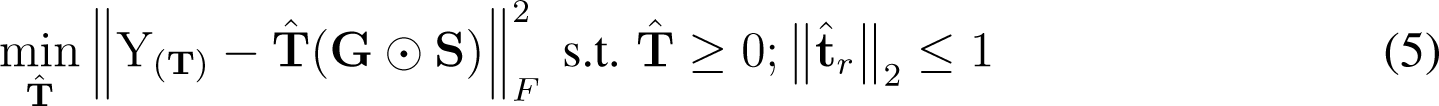

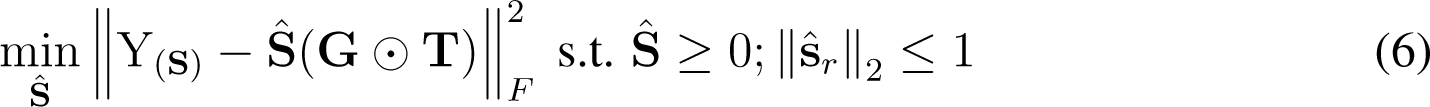

Model constraints are accommodated by modifying the inner loop least squares subproblem for a given mode. First, updates of the **G** component matrix are formulated as a lasso problem to incorporate the l1 regularization (Eq. 4). Second, updates of the **T** and **S** component matrices are achieved using a least squares problem modified to include simultaneous enforcement of the non-negativity and l2 norm constraints (Eqs. 5 and 6). Note that to make these subproblems tractable, we relax the l2 norm constraint from strict equality to an inequality, which has the effect of unit equality of the l2 norms in the sample and taxon modes when combined with the l1 penalty in the gene mode. All inner loop subproblems are solved using the Fast Iterative Shrinkage/Thresholding Algorithm (FISTA) with constraints encoded as proximal operators (*72*). To improve the convergence rate, we implemented the FISTA solver in combination with an adaptive restart scheme as proposed by O’Donoghue and Candès (2015) (*73*).

#### Implementation

Our implementation of the sparse tensor decomposition model presented here is freely available as the Python package Barnacle. Much of Barnacle’s core functionality is built on top of Tensorly (*74*). The model itself can be accessed via the ‘SparseCP’ class, modeled after Tensorly’s standardized decomposition API. Our implementation of the FISTA algorithm with adaptive restart (*72, 73*) for solving constrained inner loop least squares problems is under the ‘fista’ module of Barnacle. Code for constructing and manipulating the simulated data tensors used in model development and evaluation is available under the ‘tensors’ module. Assorted functions including visualization tools and our implementation of the cluster evaluation scores developed by Saelens et al. (2018) (*16*) are collected under the ‘utils’ module.

### Datasets

#### Simulated data

Development and evaluation of the sparse tensor decomposition model relied on simulated data. All simulations were third-order tensors constructed as the outer product of three randomly generated component matrices, constituting a signal tensor, in combination with a randomly generated Gaussian noise tensor. We varied the tensor shape, number of components, sparsity of the component matrices, and noise-to-signal ratio of each simulation depending on the experiment. Component matrix entries were randomly drawn from uniform distributions parameterized according to mode: *U* (*-*1, 1) for the simulated gene mode to allow for negative and positive values, and *U* (0, 1) for the simulated taxon and sample modes. The length of the simulated gene mode was randomly drawn from the interval (10, 1000). The lengths of the simulated taxon and sample modes were independently and randomly drawn from the interval (10, 100). The number of components (*R*) was randomly drawn from the interval (2, 10). The fraction of zero values in each component matrix was independently stipulated by drawing a value from a uniform distribution *U* (0, *m*) where 0 *≤ m ≤* 1 represents the maximum number of zeros possible so that the matrix remained full rank. Noise tensor entries were randomly drawn from a normal distribution *N* (0, 1), and then scaled them in proportion to the magnitude of the signal tensor. For each simulation, we specified the noise-to-signal ratio with a scaling factor *s*, where *s* = 10*^x^* with *x* being randomly drawn from a uniform distribution *U* (*-*1, 1). To maintain sparsity in the final simulated data tensor, we set to zero each element of the noise tensor that corresponded to a co-localized zero value in the signal tensor.

#### Metatranscriptomic data

In 2016, 2017, and 2019 a collaboration of ocean researchers collected samples for whole community metatranscriptomes (see Acknowledgments) from along a North Pacific transect ranging from around 22°N to 42°N along the 158th meridian west (fig. S1 and data S6). Replicate seawater samples were pre-filtered through either 200 micron nylon mesh (2016 samples) or 100 micron nylon mesh (2017, 2019 samples) and then filtered onto 142 millimeter polycarbonate filters and immediately flash frozen in liquid nitrogen. Samples from surface transects of all three cruises, along with the 2017 depth profiles and 20 liter incubation samples were size fractionated by serial filtration onto 3 micron and 0.2 micron polycarbonate filters, whereas the samples from the 4 liter incubations in 2017 and the 2019 depth profiles and diel study samples consisted of a single size fraction collected on 0.2 micron polycarbonate filters. Whole community RNA was extracted from each filter using the Invitrogen ToTALLY RNA extraction kit (2016 samples) and the Zymo Direct-zol RNA Miniprep Plus kit (2017, 2019 samples). To focus sequencing libraries on messenger RNA, ribosomal RNA was depleted from extractions using either the Illumina Ribo-Zero Plus kit (2016, 2017 surface samples, depth profiles, and 4 liter incubations) and the ThermoFisher RiboMinus Transcriptome Isolation kit (2019, 2017 20 liter incubations). Libraries were sequenced at the University of Washington Northwest Genomics Center using an Illumina NextSeq 500 platform (2016 samples) and an Illumina NovaSeq 6000 platform (2017, 2019 samples). We quality controlled paired sequencing reads using Trimmomatic (version 0.32), and combined reads from size fractionated samples, resulting in 222 samples of whole community RNA sequence data. Details of sample collection, RNA extraction, library preparation, and sequencing are available under the BioProject accession number associated with each dataset on the NCBI Sequence Read Archive (see Data and material availability).

We mapped sequencing reads against the 681 *Prochlorococcus* and *Synechococcus* refer-ence genomes used to generate the Cyanobacterial Cluster of Orthologous Group of proteins (CyCOG version 6.0; data S7 and S8) (*22*), using the read mapping software Salmon (version 1.10.2). We verified clade assignments for these reference genomes using a phylogeny of 424 single copy core CyCOGs that included additional genomes (data S9) (*75–77*). Single copy core CyCOGs were defined as those observed once in all 92 *Prochlorococcus* and *Synechococcus* genomes (*>* 99% complete) that were derived from cultivated isolates in CyCOG version 6.0. We used blastp (NCBI blast suite version 2.14.0) to identify CyCOGs in each additional publicly available genome. Each CyCOG protein family was aligned using clustal omega (clustalo version 1.2.4), and CyCOG protein alignments were concatenated into a single multi fasta file using MEGA (version 11.0.13). Phylogenies for the *Prochlorococcus* and *Synechococcus* concatenated protein alignments were inferred separately in FastTree (version 2.1.11) using the LG model of evolution with 100 bootstraps. We then assigned clade labels to the 681 reference genomes from CyCOG version 6.0 based on monophyletic group membership (data S7). One *Synechococcus* clade included reference genomes for strains CC9616 and KORDI-100 that have alternately been described as either clade UC-A or EnvC, which we delineate as UC-A for simplicity. Following mapping, we retained only the read counts corresponding to reference sequences annotated with both taxonomic clade and version 6.0 CyCOGs. Read counts were then aggregated by clade so that transcript abundances were consolidated to 24 pan-transcriptomes (8 *Prochlorococcus* and 16 *Synechococcus*). We restricted analysis to clades with at least one sample that recruited reads such that the percentage of detected pangenome CyCOGs surpassed 40% for *Prochlorococcus* or 60% for *Synechococcus*, corresponding to roughly 2,000 detected CyCOGs. The following clades met this threshold and were retained for further analysis: *Prochlorococcus* HLI, HLII, and LLI, and *Synechococcus* I, II, III, IV, CRD1, CRD2, and UC-A.

#### Normalization

To disentangle transcript abundance from organism abundance and focus our analysis on the most differentially expressed genes, we used version 2 of the variance stabilizing transform (vst) (*24*) to normalize the mapped read count matrix of each clade. Briefly, the vst fits a separate generalized linear model to the transcript abundance counts of each gene, using a negative binomial distribution as the link function (*23*), which accommodates the overdispersed mean-variance relationship observed in raw transcript abundance profiles (fig. S8). In each sample the transcript abundance of the gene is modeled as a function of overall clade abundance, approximated by the the total number of reads recruited to the clade pangenome in that sample. Regression parameters are then regularized across all gene models within each clade pangenome. Model residuals are output as the normalization product, indicating the degree and direction to which the transcript reads detected in each sample diverge from the expectation of the vst model. Before applying the normalization method, we pre-processed the read count matrix of each clade with detection and coverage thresholds. These thresholds retained genes detected in at least 3 samples, and samples in which the percentage of detected genes achieved at least 1% of the maximum per-sample coverage observed in the clade pangenome. To account for dataset-scale bias in transcript counts that arise from technical artifacts (e.g. discrepancies in sample processing, library preparation, sequencing platform, etc.) we passed the sequencing batch to the vst function to be regressed out as a nuisance variable. All other vst arguments were set to the version 2 defaults, except ‘min cells’, which was set to match the 3 sample detection threshold. This procedure produced for each clade a matrix of normalized residual transcript abundance values in which variance was de-correlated from mean abundance (fig. S9).

#### Data tensorization

The normalized residual transcript abundance matrices were segregated by genus and arranged into two gene (CyCOG) *⇥* taxon (clade) *⇥* sample tensors by aligning CyCOGs between clades. Any CyCOGs or samples not detected in a particular clade were filled in with zero values. The resulting *Prochlorococcus* data tensor encompassed 5,084 CyCOGs, 3 clades (HLI, HLII, and LLI) and 178 samples (data S10). The *Synechococcus* data tensor encompassed 6,161 CyCOGS, 7 clades (I, II, III, IV, CRD1, CRD2, and UC-A), and 222 samples (data S11).

### Metrics

The metrics used to assess model performance are summarized in Table 2. The sum of squared errors (SSE) quantified model fit, and was calculated using the ‘relative error’ function of the ‘tlviz’ Python package for the analysis and visualization of tensor decompositions (*78*). SSE was calculated relative to the Froebenius norm of each data tensor so that scores could be compared across datasets. The factor match score (FMS) quantified the similarity of two sets of component matrices, and was calculated using the ‘factor match score’ method of the ‘tlviz’ package. Importantly, the order of components in a tensor decomposition model is arbitrary, so prior to comparing models, the order of components was permuted against a reference to align the closest matching components between models. This was accomplished using the ‘permute components’ function of the ‘tlviz’ package, which employs the ‘linear sum assignment’ function of ‘scipy.optimize’.

Precision, recall, and F1 metrics were used to assess the accuracy of gene clusters derived from model components. Both precision and recall measure the overlap in gene membership between model and ground truth clusters. Precision measured the overlap as a proportion of model cluster membership, and recall as a proportion of ground truth cluster membership. By design, genes are permitted to be included in more than one of the clusters derived from model gene component matrices. We therefore selected implementations of precision and recall metrics proposed by Saelens et al. (2018) (*16*), which accommodate clusters with overlapping gene membership. For a balanced measure of cluster accuracy, the F1 score was calculated as the harmonic mean of the Saelens et al. precision and recall scores.

### Model fitting

All instantiations of sparse tensor decomposition models were fit using the ‘SparseCP’ API of the Barnacle package. We oriented data tensors gene *⇥* taxon *⇥* sample so that the l1 sparsity penalty was applied to the first mode, and non-negativity and unit l2 norm constraints were applied to the second and third modes. In cases where the sparsity coefficient was set to *.λ* = 0, the unit l2 norm constraint was removed from all modes.

Convergence was defined using loss, calculated as

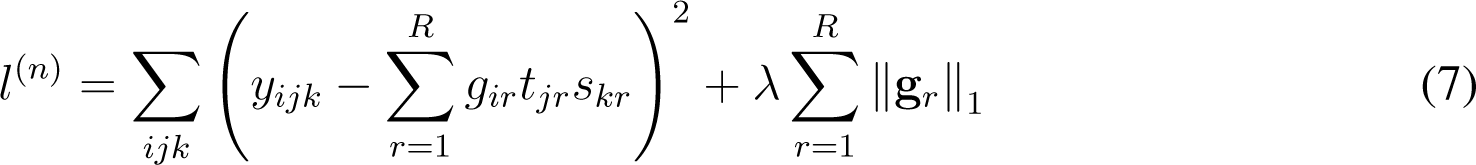

where *l*^(^*^n^*^)^ designates the loss at iteration *n* of the modified ALS algorithm. In all experiments performed in this study, we considered the algorithm to have converged when the change in loss dropped below 10*^-^*^5^, or equivalently, when the following inequality was satisfied:

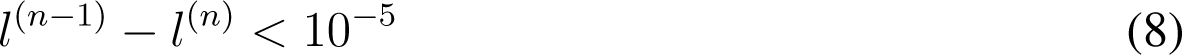

This change in loss is always non-negative because the loss decreases monotonically with each iteration of the ALS algorithm.

Depending on the initialization, it is possible for the ALS algorithm to converge on a local minimum rather than the globally optimal solution (*15*). To mitigate this issue, we repeated all decompositions with five different random initializations, and the solution corresponding to the lowest loss was saved for analysis. Convergence was verified for each initialization based on the loss criterion outlined (Eqs. 7 and 8); all decompositions performed in this study converged in under 2,000 iterations. The random state of each decomposition was initialized with a unique integer seed, and to ensure reproducibility the seed of each saved model was stored as a local text file alongside model solutions and parameters.

#### Parameter selection

To select appropriate values of *R* and *.λ*, we developed a cross-validated grid search strategy that made use of sample replicates in the metatranscriptomic datasets. Data tensors were split by replicate along the sample axis to produce three replicate subtensors of shape *I ⇥ J ⇥ K_A_*, *I ⇥ J ⇥ K_B_*and *I ⇥ J ⇥ K_C_*, where *I* is the number of genes in the full dataset, *J* is the number of taxa in the full dataset, and *K_A_*, *K_B_* and *K_C_* are the number of samples in replicate set A, B and C, respectively. For experiments with simulated data, we simulated replicates by generating three independent noise tensors, and combining each with the same underlying signal tensor. For both the simulation and real data tensors, we fit a series of models to each replicate subtensor using a grid search of different *R* and *.λ* parameter values. Six cross-validated SSE scores were calculated for each unique set of parameters by comparing each fit model against the two held out replicate subtensors. Three cross-validated FMS scores were calculated for each parameter set by comparing the components between each pair of replicate models. In the simulated data experiments, every combination of parameters *R 2* [1, 2, 3, *…,* 12] and *.λ 2* [0.0, 0.05, 0.1, 0.2, 0.4, 0.8, 1.6, 3.2, 6.4, 12.8] was tested against each simulation. In fitting both the *Prochlorococcus* and *Synechococcus* data tensors, we first pursued a coarse parameter grid search of *R 2* [1, 5, 10, 15, 20, 25, 30, 35, 40, 45, 50] and *.λ 2* [0.0, 0.1, 1.0, 10.0, 100.0] in order to explore general trends. We examined the cross-validated SSE scores of the *.λ* = 0.0 models to identify the the minimum error model in the absence of l1 regularization. The number of components was set to the *R* value corresponding to the this minimum. After a fine-tuned search of sparsity coefficients with *.λ 2* [1., 2., 3*., …,* 32., 64.] we then selected the sparsity coefficient as the maximum *.λ* value at which the cross-validated FMS remained within one standard error of the maximum FMS.

#### Bootstrapping

For the *Prochlorococcus* and *Synechococcus* models, we used bootstrapping in conjunction with cross validation in order to mitigate bias originating from inconsistent numbers of replicates between samples (fig. S1), and to estimate the consistency of the resulting models. The 222 metatranscriptome samples originate from 87 unique conditions, of which 55 were sampled in triplicate, 25 in duplicate, and 7 were represented by a single sample. In each bootstrap, ‘A’, ‘B’, and ‘C’ replicate labels were randomly shuffled for each of the 55 conditions sampled in triplicate. For conditions with less than three replicates, replicate labels were randomly drawn without replacement. This ensured that the three resulting replicate subtensors were roughly equivalent in size, and that every sample had an equal and independent chance of being placed in any replicate set in any given bootstrap. Models were then fit independently to each replicate subtensor. Cross-validated SSE and FMS scores were calculated using only the indices of the samples common to the pair of replicate subtensors being compared. Consequently, conditions represented by a single replicate were used in fitting the models, but were not included in score calculations. For every unique set of parameters in the grid search, we fit 10 bootstraps, each with three replicate subtensors, for a total of 30 randomly shuffled data subtensors. For the best fit models, we fit 100 bootstraps of 3 replicate subtensors each, for a total of 300 shuffled replicate subtensors.

### Model evaluation and interpretation

#### Robustness to mis-specification

We evaluated the effect of parameter mis-specification on model performance using simulated data, and these experiments and results are described in the Supplementary Text. We also assessed the robustness of model components to parameter mis-specification in the real metatranscriptomic datasets. Among bootstrap models parameterized with best fit parameters (as determined by cross-validated grid search), a “best representative” model was identified as the bootstrap with the highest mean FMS, calculated in comparison to all other bootstrap models with best fit parameters. Models fit with alternate, suboptimal *R* and *.λ* parameters were then aligned to this best representative model. Next, individual components were extracted from each aligned model and component-specific FMS scores were assessed for each component by comparing it against its matching component in the aligned best representative reference model. Additionally, the percentage of variation represented by each component of the best fit models was calculated using the ‘percentage variation’ function of the ‘tlviz’ package (*78*).

#### Comparison of component weight profiles

For each component, we compiled a consensus weight profile from the collection of model bootstraps. Gene, taxon, and sample weight vectors were collected from each of 300 bootstrap replicates of best fit models, aligned against the best representative model. In the gene-mode, the median gene weight was calculated for each CyCOG, and the collection of CyCOGs with non-zero median weights was designated as a cluster. Additionally, the proportion of bootstraps in which the gene weights of a given CyCOG were non-zero was calculated as a measure of the strength of the association of a given CyCOG to a given cluster. The full set of *Prochlorococcus* component weight profiles can be found in Supplementary Dataset 1, and *Synechococcus* component weight profiles in Supplementary Dataset 2.

We compared gene-, taxon-, and sample-mode weight profiles between combined *Prochlorococcus* and *Synechococcus* components, considering only the most robust components with a cross-validated component FMS of 0.5 or higher. This threshold removed components pro1, pro14, syn1, syn2, and syn13 from further analysis. For each mode, a Pearson correlation was calculated between the median weight vectors of each pair of components, and the resulting correlation matrix was hierarchically clustered to identify groups of components with similar weight profiles. Additionally, a hierarchically clustered correlation matrix was calculated for a composite weight profile constructed by concatenating the median weight vectors of the gene-, taxon-, and sample- modes. To enable comparisons between *Prochlorococcus* and *Synechococcus* model components, any samples and CyCOGs specific to one model were added to the other model and filled in with zero values.

We also compared clusters from our analysis against previously identified clusters, published in a study that examined diel gene expression of *Prochlorococcus* strain MED4 grown over a light/dark cycle under nutrient replete laboratory conditions (Zinser et al., 2009) (*29*). We used blastp (NCBI blast suite version 2.14.0) to identify the CyCOG associated with each MED4 gene (data S4), so that the CyCOG content of the Barnacle clusters could be directly compared to the CyCOG content of the Zinser clusters. Zinser et al. used a soft clustering algorithm to determined gene cluster membership, in which a score between 0 and 1 quantified the strength of the association of each gene to its cluster assignment, similar to the percent bootstraps score calculated in this study. We used the ‘f1 score’ function of the ‘sklearn.metrics’ module to calculate a weighted F1 score between each pair of Zinser and Barnacle clusters, with weights set to the sum of Zinser cluster scores and Barnacle percent bootstrap scores. Null F1 scores were generated from randomly shuffled Zinser and Barnacle weight vectors, and an empirical cumulative density function (ECDF) constructed from the null scores was used to identify significantly similar cluster pairs, using a one-sided p-value threshold of 0.05 and Benjamini-Hochburg adjustment to control false discovery rate with multiple comparisons. We used the Python library ‘statsmodels’ to calculate the ECDF and adjust p-values, employing the ‘ECDF’ function of the ‘statsmodels.distributions.empirical distribution’ module and the ‘multipletests’ function of the ‘statsmodels.stats.multitest’ module, respectively.

We compared component sample weight profiles against measurements of environmental conditions, accessed via the Simons Collaborative Marine Atlas Project (Simons CMAP) (*79*). Estimates of surface chlorophyll were taken from the MODIS chlorophyll dataset in CMAP, an 8-day averaged product calculated using sea surface measurements from the MODIS Aqua satellite. The University of Southern California Marine Trace Elements Lab contributed dissolved iron measurements, and the Dave Karl lab at the University of Hawaii – Manoa contributed dissolved nitrate plus nitrite measurements.

#### Inference of circadian expression peak

We used a weighted kernel density estimate (KDE) to estimate the peak expression time of each component profile, assuming a circadian expression pattern. Across the combined metatranscriptome datasets, sampling times were not uniformly distributed around the 24 hour cycle. To account for uneven sample times, we fit a baseline KDE to the sample times of each of the *Prochlorococcus* and *Synechococcus* datasets, assigning equal weight to every sample. Then for each cluster, we fit a second weighted KDE to the component sample times, using as weights the product of the median sample-mode weight profile and the inverse of the appropriate baseline KDE. In effect, this biases the KDE towards samples in which the component signal is highly expressed while counteracting the KDE’s inherent bias towards densely sampled times of day. The maximum of the weighted KDE was taken as the estimated peak expression time of each component. We used the ‘gaussian kde’ function of the ‘scipy.stats’ Python library to fit KDEs.

#### Functional enrichment of gene clusters

We analyzed the biological function of each cluster by testing for over-representation of genes involved in common metabolic pathways or cell processes cataloged in the KEGG database (release 109.0). First, a consensus functional annotation was assigned to each CyCOG based on the best scoring annotation of a majority of member genes. Most CyCOGs had no annotation assigned to any member gene (36,693/40,295; 91.06%), but among the annotated CyCOGs, the majority had concordant functional annotations among member genes (3,474/3,602; 96.44%). Next, the majority annotations were used to assign CyCOGs to a subset of KEGG pathways that we manually curated to remove irrelevant processes, such as human disease pathways. Finally, the enrichment of each KEGG pathway was evaluated for each Barnacle cluster using a one-sided Mann-Whitney U test, as implemented by the ‘mannwhitneyu’ function of the ‘scipy.stats’ library. For each cluster, the test compares the gene weights of CyCOGs belonging to a particular pathway to the gene weights of CyCOGs not in that pathway, and evaluates the null hypothesis that the distribution of pathway gene weights is less than or equal to the distribution of non-pathway gene weights. In effect, this identifies the pathways for which member CyCOGs are significantly over-represented in the cluster, as compared to their background frequency in the expressed pangenome. We delineated enriched KEGG pathways as those with a p-value less than 0.01, following Benjamini-Hochburg adjustment for multiple comparisons using the ‘statsmodels.stats.multitest’ library. Consensus CyCOG annotations and KEGG pathway enrichments for each cluster can be found in Supplementary Dataset 3.

## Acknowledgments

We thank the scientists and crew who supported the SCOPE Gradients expeditions, especially Randie Bundy, Bryndan Durham, Ryan Groussman, Nicholas Hawco, Rhonda Morales, and Megan Schatz for collecting and processing samples for metatranscriptome sequencing. In addition, thank you to Yngve Mardal Moe for help with the FISTA implementation, Chris Berthiaume for help with parallelization, James Mullet for help updating reference genome clade assignments, and all the members of the Armbrust and Noble labs for insights, discussions, and feedback.

## Author Contributions

Conceptualization, Formal Analysis, Visualization, and Writing – Original Draft: S.B. Methodology and Software: S.B., M.R. Data Curation: S.B., P.M.B. Supervision and Funding Acquisition: E.V.A. Writing – Review and Editing: S.B., M.R., P.M.B., R.B., E.V.A.

## Data and Materials Availability

All data needed to evaluate the conclusions in the paper are present in the paper and/or the Supplementary Materials. Supplementary Data files are available in an associated Zenodo repository: https://doi.org/10.5281/zenodo.12210994. The sequences reported have been deposited in the NCBI Sequence Read Archive under the following BioProject accession numbers: PRJNA1088528, PRJNA1090042, PRJNA1090086, PRJNA1090467, PRJNA1090899, PRJNA1091352, PRJNA492143, and PRJNA816919. Source code and documentation for the Barnacle model can be found in the package repository: https://github.com/blasks/barnacle.

All code required to process data, run analyses and produce figures presented in this manuscript can be found in an associated manuscript repository: https://github.com/blasks/barnacle-manuscript.

