## Supplementary Materials for "Simultaneous acclimation to nitrogen and iron scarcity in open ocean cyanobacteria revealed by sparse tensor decomposition of metatranscriptomes"

Stephen Blaskowski *et al.*

**This PDF file includes:**

Supplementary Text
Figs. S1 to S16
Legends for data S1 to S11

**Other Supplementary Materials for this manuscript include the following:**

Data S1 to S11

### Supplementary Text

Validation of parameter selection strategy with simulated data

We developed a cross-validated grid search strategy that leveraged sample replicates to select best fit model parameters (See Materials & Methods). We evaluated the effectiveness of the strategy using 100 simulated tensors that collectively represented a range of tensor shapes, ranks, sparsity patterns, and noise-to-signal ratios ranging from 0.1 to 10. We simulated sample replicates by generating three identical copies of each signal tensor and combining each with an independent Gaussian noise tensor. We then independently fit models to each replicate tensor, calculated cross-validated SSE and FMS scores by comparing between replicates, and selected parameters of best fit. We selected the best fit $R$ parameter as the number of components that resulted in the lowest cross-validated SSE, and among models fit with this value of $R$, we selected the best fit sparsity coefficient as the maximum $\lambda$ value at which the cross-validated FMS fell within one standard error of the maximum FMS.

We compared $R$ and $\lambda$ parameters selected via cross-validated grid search with ground truth parameters, as determined by the true number of components used to generate each simulation and the value of $\lambda$ that resulted in the maximum F1 score (fig. S3). The $R$ identified by the grid search matched the true number of components in 86 of 100 simulations and was off by no more than 1 component in 92 simulations (fig. S4A). When the noise-to-signal ratio was greater than 1, the selected $R$ in 13 simulations differed from the true number of components by an average of 2.2. At noise-to-signal ratios below this threshold, the selected $R$ in a single simulation differed by one from the true number of components. The selected $\lambda$ matched the optimal $\lambda$ in 46 simulations and was within a twofold change in 80 simulations (fig. S4B). The frequency and magnitude of $\lambda$ mis-specification was relatively consistent across simulation noise levels. Underestimation was more common than overestimation for both $R$ and $\lambda$: 11 of the 14 $R$ mis-specifications and 34 of the 54 $\lambda$ mis-specifications undershot the ground truth parameter value. These results demonstrated that the cross-validated grid search strategy selects parameters that approach optimal values, even up to a noise level ten times that of the signal.

We also examined the ramifications of inaccurate parameter selection in models fit to the 100 simulated data tensors. We aligned all models against the ground truth components used to generate each simulation, and assessed SSE, FMS, precision, recall, and F1 scores in reference to a noiseless version of the simulated dataset. In most cases, the FMS, precision, recall, and F1 scores showed little difference between models with mis-specified or true $R$ parameter values (fig. S5, A to D). In all but one case, the F1 score of models parameterized with an inaccurate $R$ was nearly identical to the optimally-parameterized model, highlighting that model-derived clusters are robust to $R$ mis-specification. When $\lambda$ was underestimated, the resulting gene clusters generally exhibited lower precision and higher recall, whereas when $\lambda$ was overestimated, the clusters exhibited higher precision and lower recall (fig. S5, E to H). These data suggested that although high-sparsity models may incorrectly exclude some genes from clusters, increasing sparsity provides greater confidence in the retained composition of modules, that they accurately reflect true cluster structure in the underlying data.

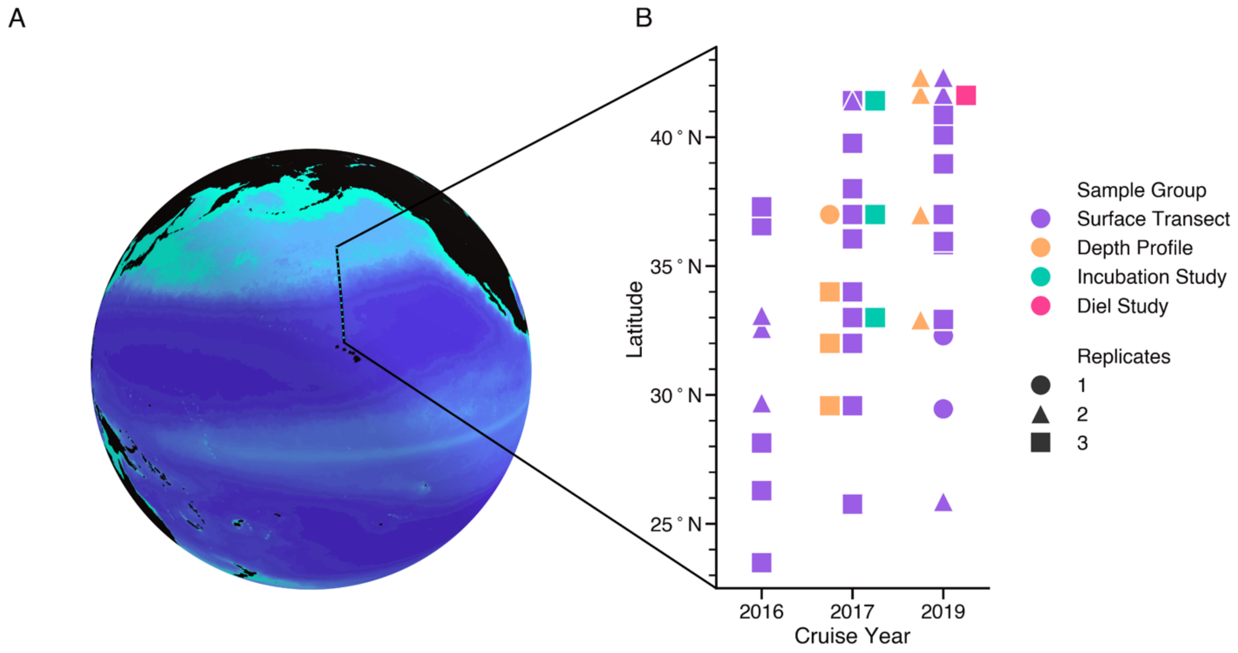


**Fig. S1. Summary of sampling locations and datasets integrated in this study.**
(A) Approximate cruise track (dotted line) along 158th meridian west, plotted over April climatology of surface chlorophyll, as measured by the MODIS Aqua satellite. (B) Sampling latitudes by cruise year, colored by dataset. Marker shape indicates maximum number of replicates retrieved for each sample set. Each marker for a depth profile, incubation study, or diel study encompasses multiple samples taken at the same latitude at different depths, treatments, and times of day, respectively. See Supplementary data 6 for detailed sample metadata.

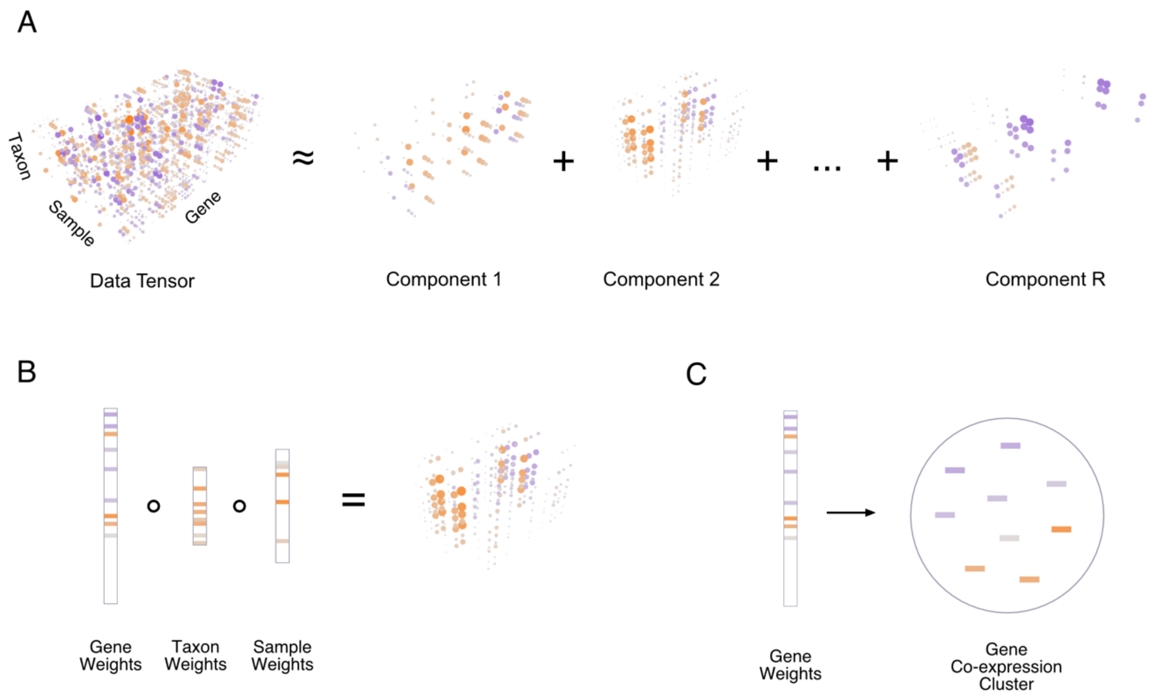


**Fig. S2. Diagram of sparse tensor decomposition model**

(A) Visual representation of sparse CP tensor decomposition in which data tensor is modeled as a sum of *R* sparse components. Orange indicates positive values, purple negative values, and marker size indicates the magnitude of the value. (B) Diagram of a single component, which consists of gene, taxon, and sample weights. The outer product of these weight vectors equals a rank-1 tensor that represents the transcript abundance pattern modeled by the component. (C) Diagram illustrating the derivation of a gene co-expression cluster from the genes corresponding to non-zero weights in a component gene weight vector.

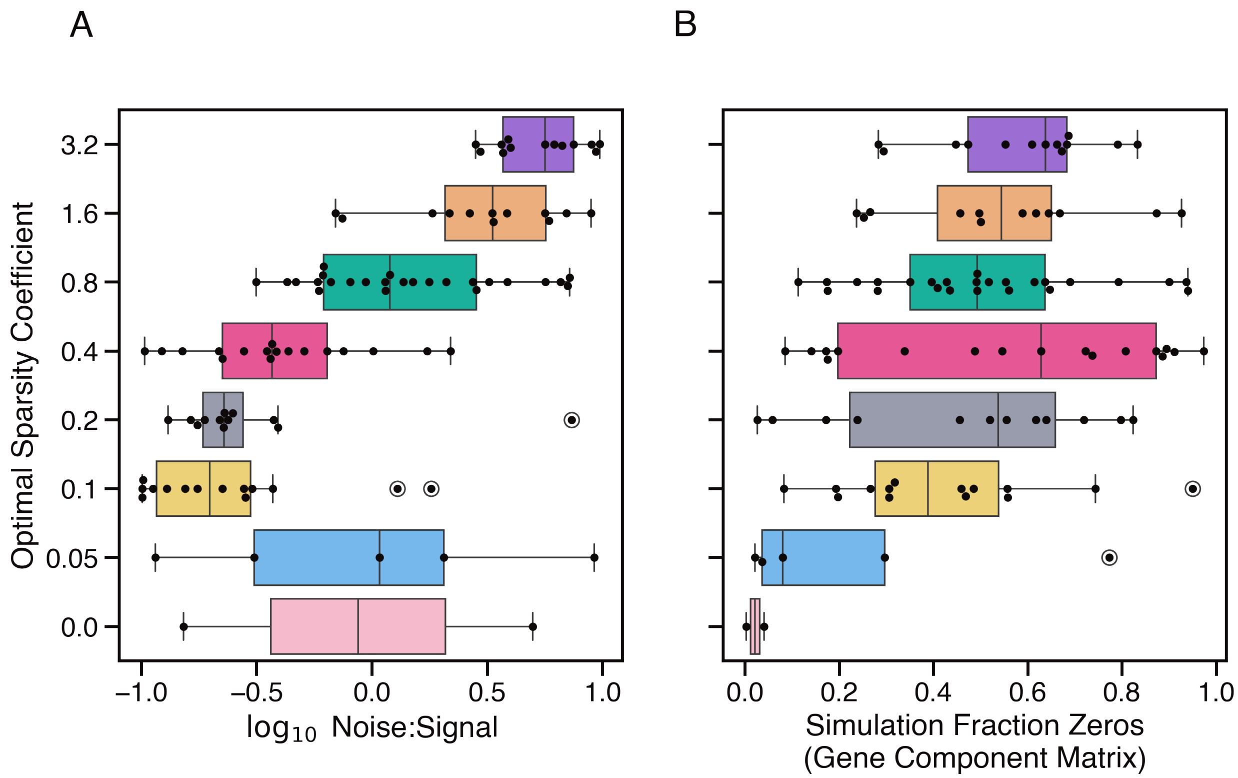


**Fig. S3. Relationship of estimated optimal sparsity coefficient to simulation noise and sparsity of the gene component matrix.**

Optimal sparsity coefficient (λ) of models fit to 100 simulated datasets, plotted as a function of (A) simulated noise-to-signal ratio and (B) fraction of zero values in the gene component matrix used to generate the simulation. Points are binned by optimal sparsity coefficient, estimated via parameter grid search as the λ corresponding to the maximum F1 score between model and simulation gene clusters. Boxes show inner 50th percentile of bin distributions, centered on the median, whiskers delineate range, and circles show outlier points excluded from distribution calculations.

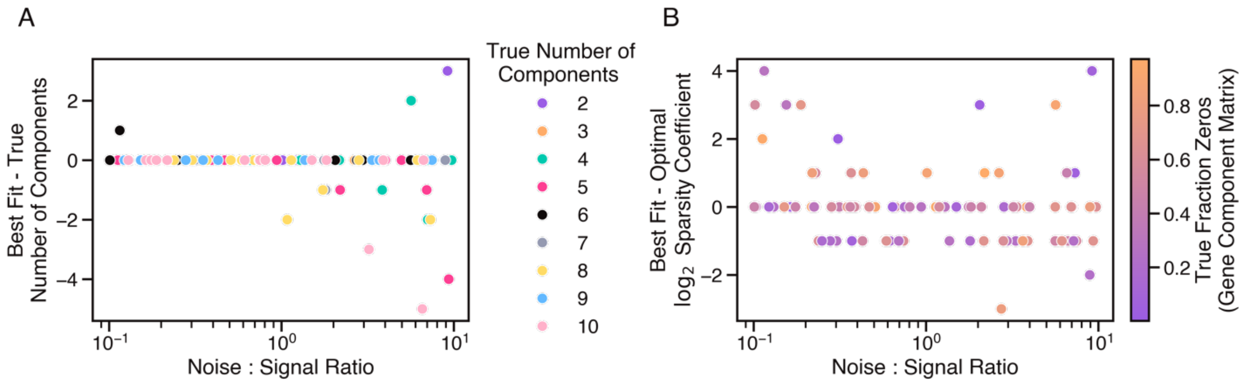


**Fig. S4. Cross-validated grid search identifies best fit parameters that approach ground truth in 100 simulated datasets.**

(A) Difference between model *R* parameter (number of components) selected via cross-validated grid search, and the true number of components in each simulation, plotted against simulation noise level and colored by true *R*. (B) Log-2 change between λ parameter (sparsity coefficient) selected via cross-validated grid search, and the optimal λ that maximizes the mean F1 score between model-derived gene clusters and those of the simulation ground truth. Scores are plotted against simulation noise level and colored by the true fraction of zero values in the simulation gene component matrix. Six data points in which the optimal or best fit λ was 0 were excluded for ease of visualization.

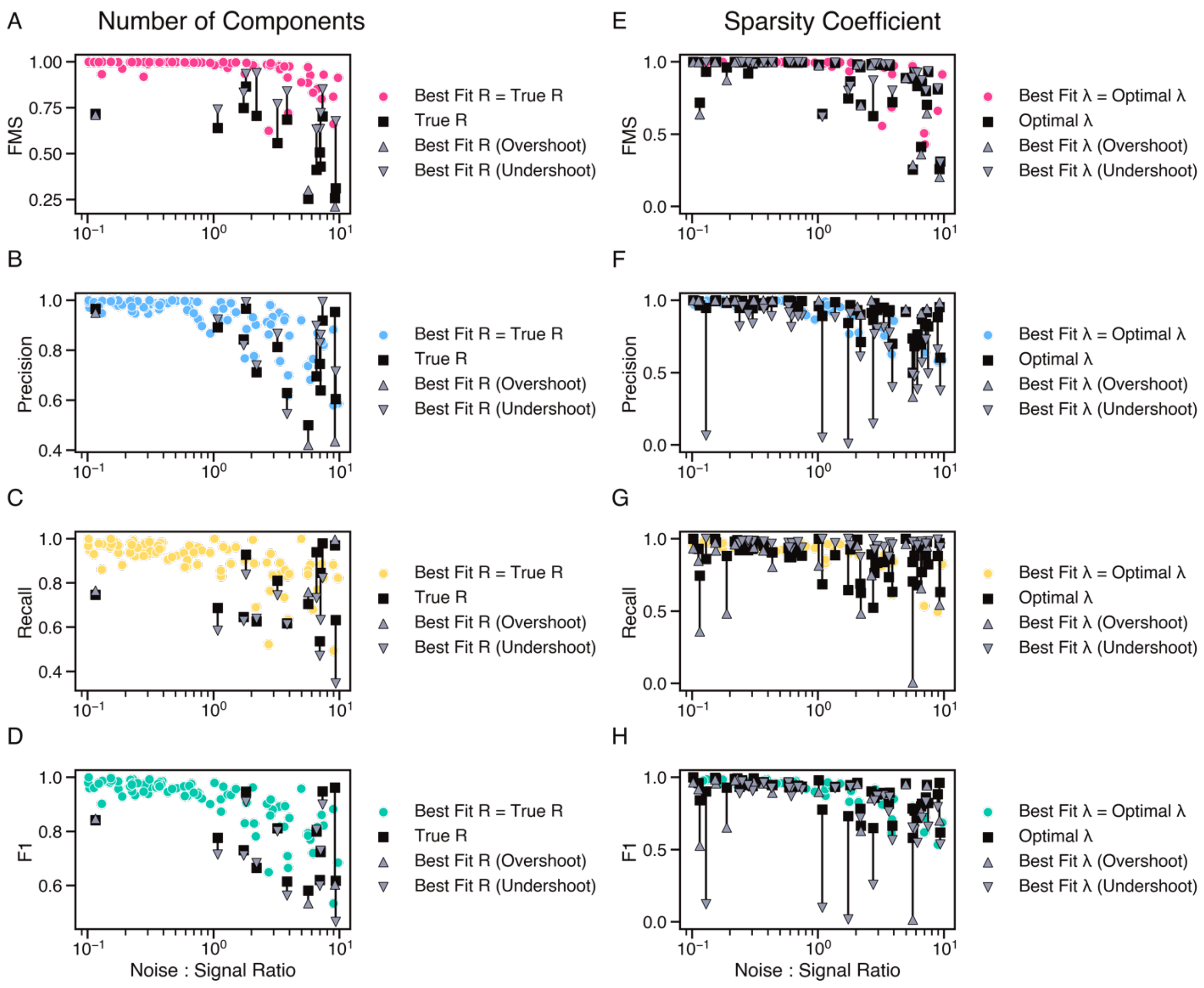


**Fig. S5. Evaluation of effect of parameter mis-specification on model component accuracy in 100 simulated datasets.**

The effect of *R* parameter (number of components) mis-specification on (A) overall model factor match score (FMS), (B) gene cluster precision, (C) recall, and (D) F1 score, evaluated in comparison to simulation ground truth. The effect of λ parameter (sparsity coefficient) mis-specification on (E) overall model FMS, (F) gene cluster precision, (G) recall, and (H) F1 score, evaluated in comparison to simulation ground truth. Colored points indicate simulations in which cross-validated grid search identified the optimal parameter. In cases where cross-validation undershot or overshot the optimal parameter, black bars indicate the difference in score between model fit with mis-specified parameter (grey triangles) and optimal parameter (black square).

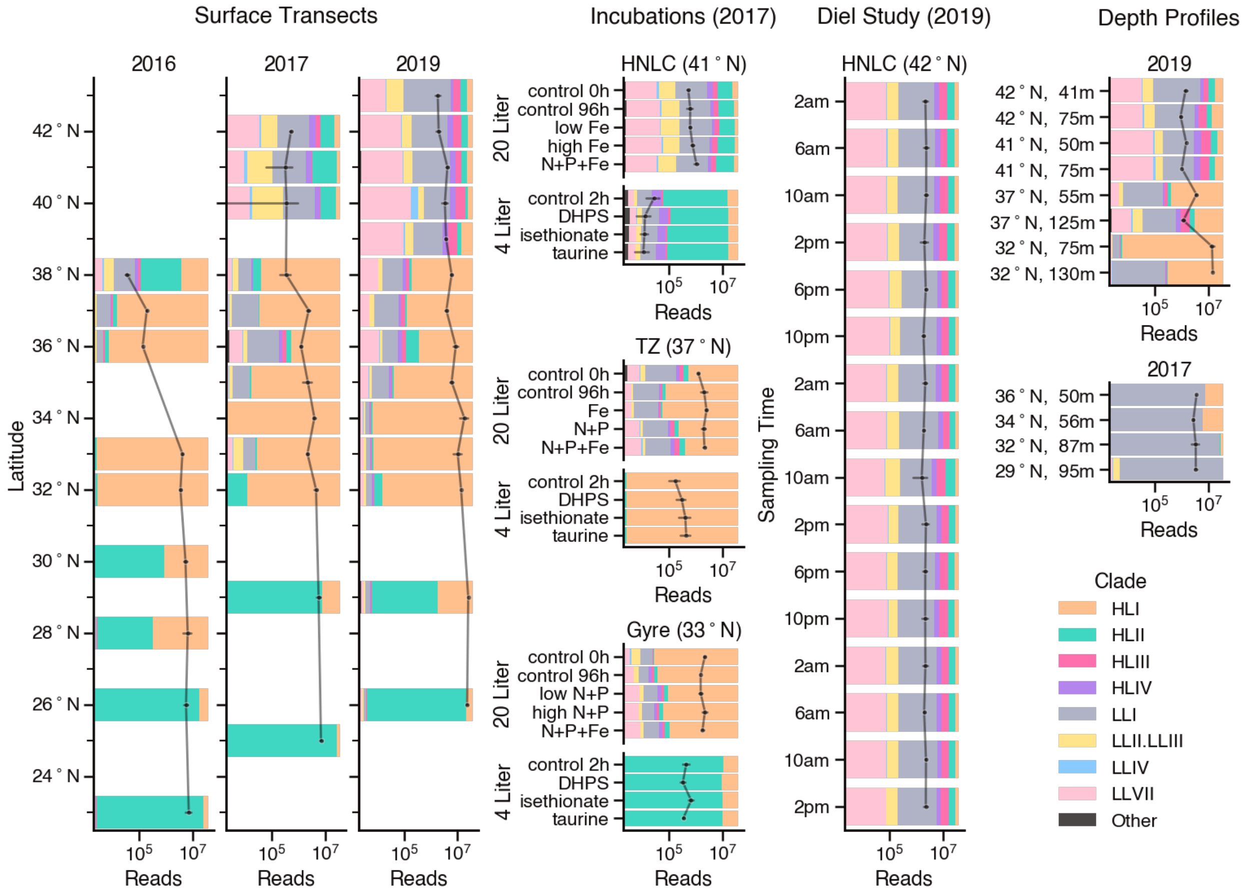


**Fig. S6. Composition of *Prochlorococcus* community transcript sequencing reads.** Relative abundance of transcript reads mapped to *Prochlorococcus* clades. The size of each colored region is proportional to the fraction of sample reads mapped to the indicated clade (sample replicates averaged) with clade fractions summing to one in each sample, and samples grouped by dataset. X-axis corresponds to black line, showing aggregate count of *Prochlorococcus* transcript reads per sample with error bars indicating standard deviation of replicates.

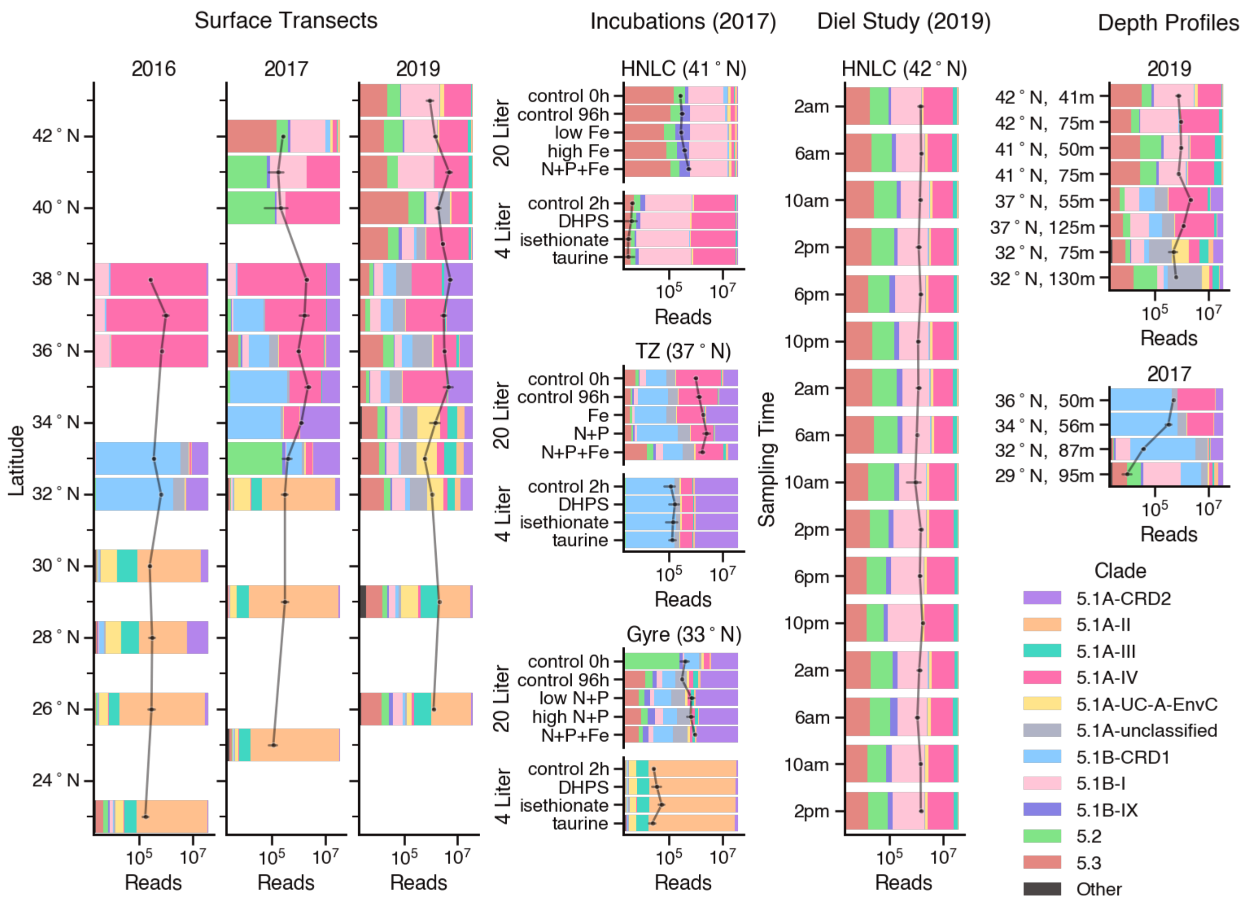


**Fig. S7. Composition of *Synechococcus* community transcript sequencing reads.** Relative abundance of transcript reads mapped to *Synechococcus* clades. The size of each colored region is proportional to the fraction of sample reads mapped to the indicated clade (sample replicates averaged) with clade fractions summing to one in each sample, and samples grouped by dataset. X-axis corresponds to black line, showing aggregate count of *Synechococcus* transcript reads per sample with error bars indicating standard deviation of replicates.

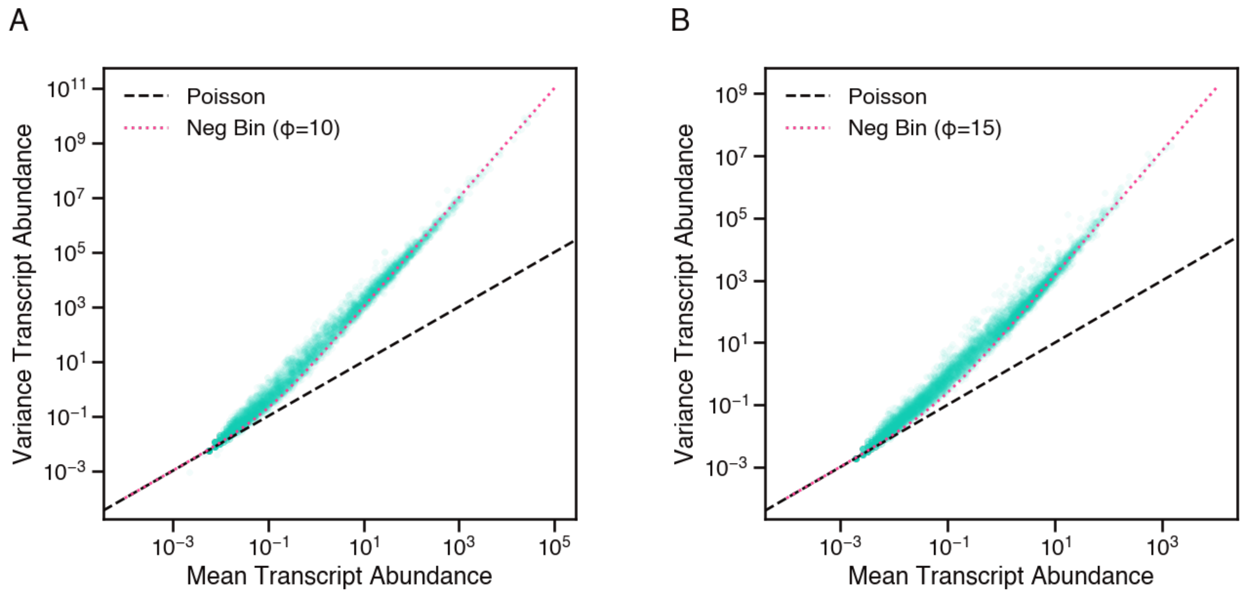


**Fig. S8. Raw transcript abundance counts exhibit an overdispersed mean-variance relationship.**

Variance ($\sigma^2$) vs. mean ($\mu$) of (A) *Prochlorococcus* and (B) *Synechococcus* transcript abundance counts. Each point represents the transcript abundance profile of an individual gene (CyCOG) across samples mapped. As a visual guide, the mean-variance relationship of a Poisson distribution ($\sigma^2 = \mu$) is plotted as a dashed black line, and the mean-variance relationship of a negative binomial distribution ($\sigma^2 = \mu + \frac{\mu^2}{\phi}$) is plotted as dotted red line, parameterized with an appropriate overdispersion parameter, $\phi$.

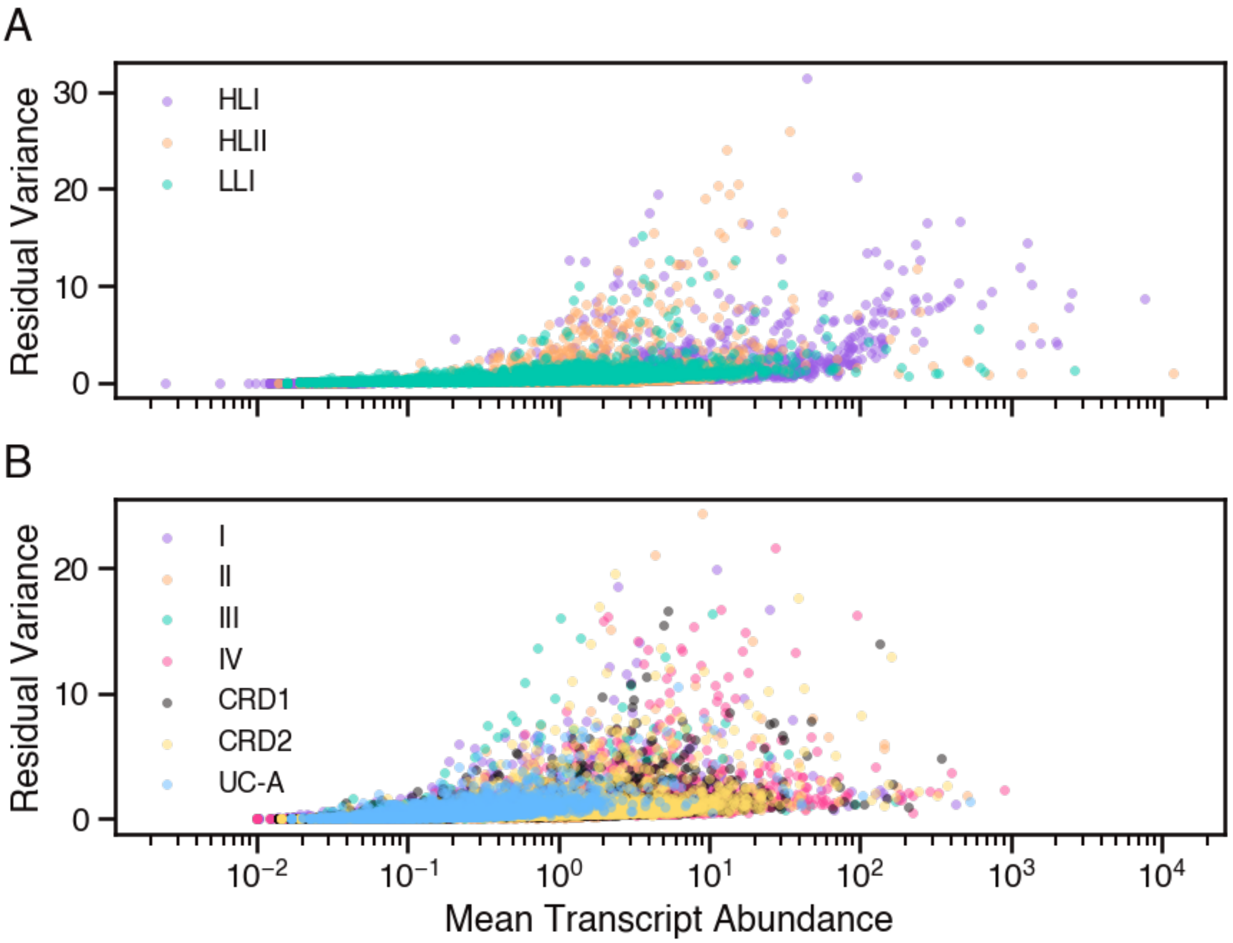


**Fig. S9. Variance is de-correlated from mean in normalized residual transcript abundance values.**

Variance of normalized residual transcript abundance vs. mean raw transcript abundance of (A) *Prochlorococcus* and (B) *Synechococcus* data. Each point represents an individual gene (CyCOG), and points are colored by clade of origin.

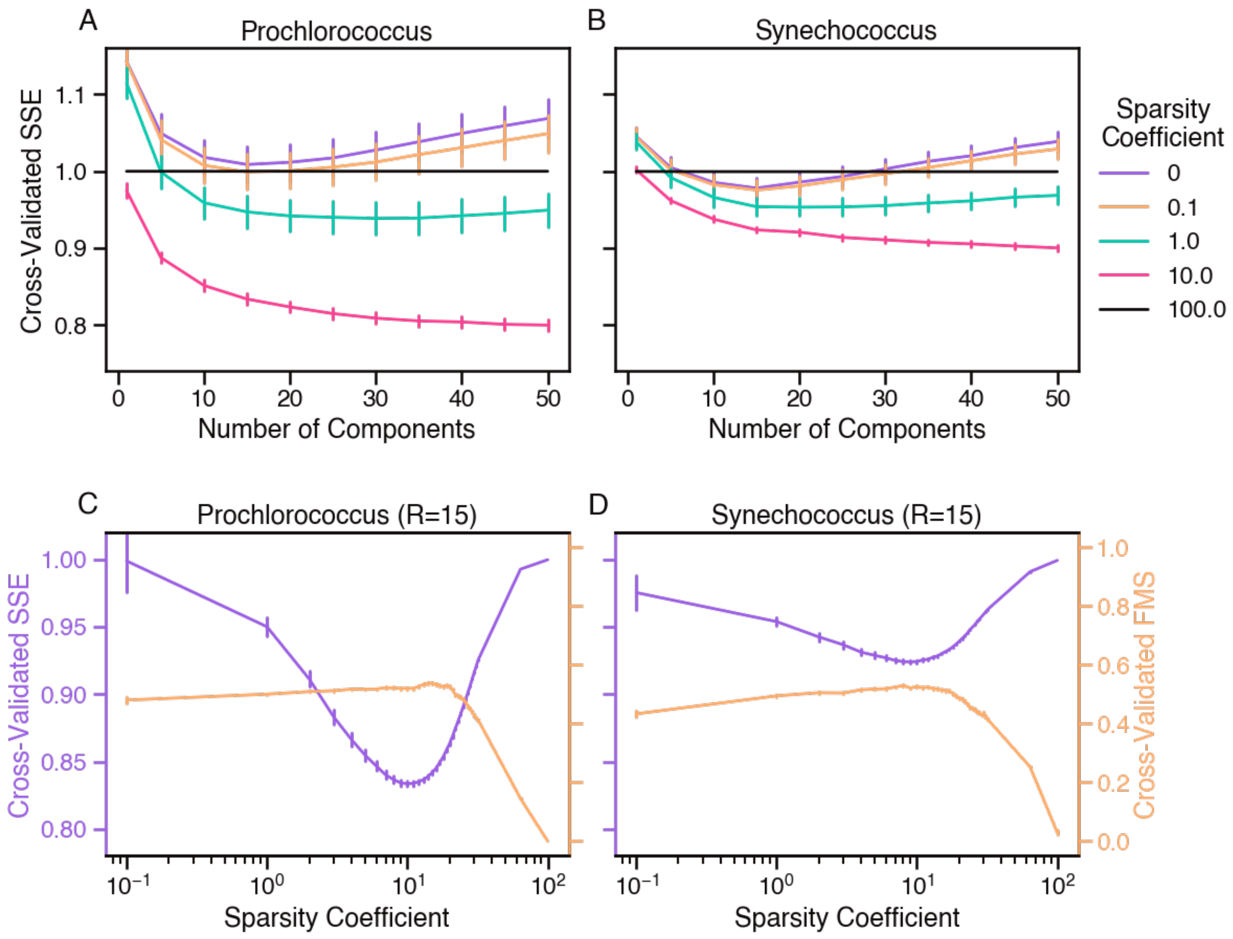


**Fig. S10. Results of cross-validated parameter grid search fitting model to *Prochlorococcus* and *Synechococcus* residual transcript abundance data.**

(A,B) Cross-validated sum of squared errors (SSE) of sparse tensor decomposition models parameterized with different numbers of components ($R$) and sparsity coefficients ($\lambda$) and fit to (A) *Prochlorococcus* and (B) *Synechococcus* residual transcript abundance data. (C,D) Cross-validated SSE and factor match score (FMS) as a function of sparsity coefficient ($\lambda$) of (C) *Prochlorococcus* models and (D) *Synechococcus* models fit with $R = 15$ components. Each data point is the mean of 10 bootstrapped datasets, and bars show standard error of the mean.

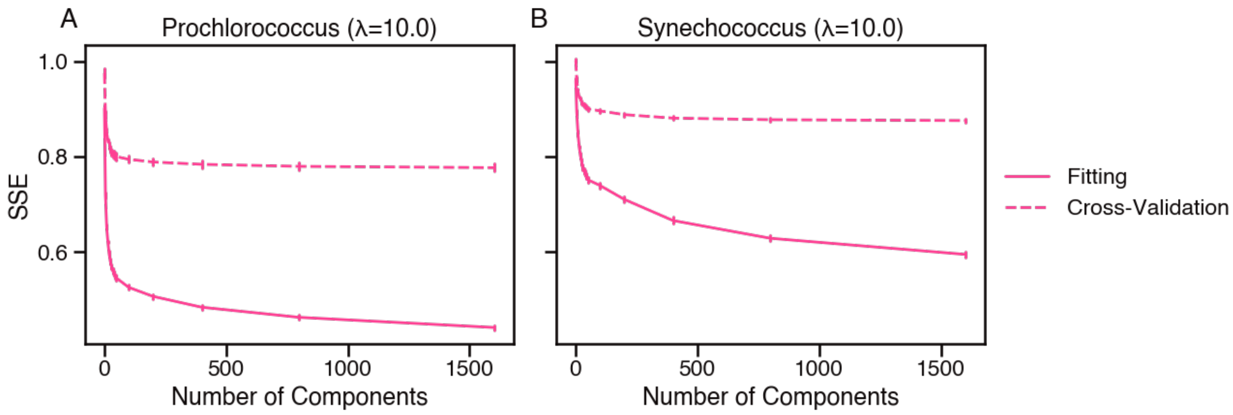


**Fig. S11. SSE monotonically decreases with an increasing number of components up to *R* = 1600 in models with λ = 10.0.**

Fitting and cross-validated sum of squared errors (SSE) of (A) *Prochlorococcus* and (B) *Synechococcus* models parameterized with a sparsity coefficient of λ = 10.0 and different numbers of components ranging from $R = 1$ to $R = 1600$. Each data point is the mean of 10 bootstrapped datasets, and bars show standard error of the mean.

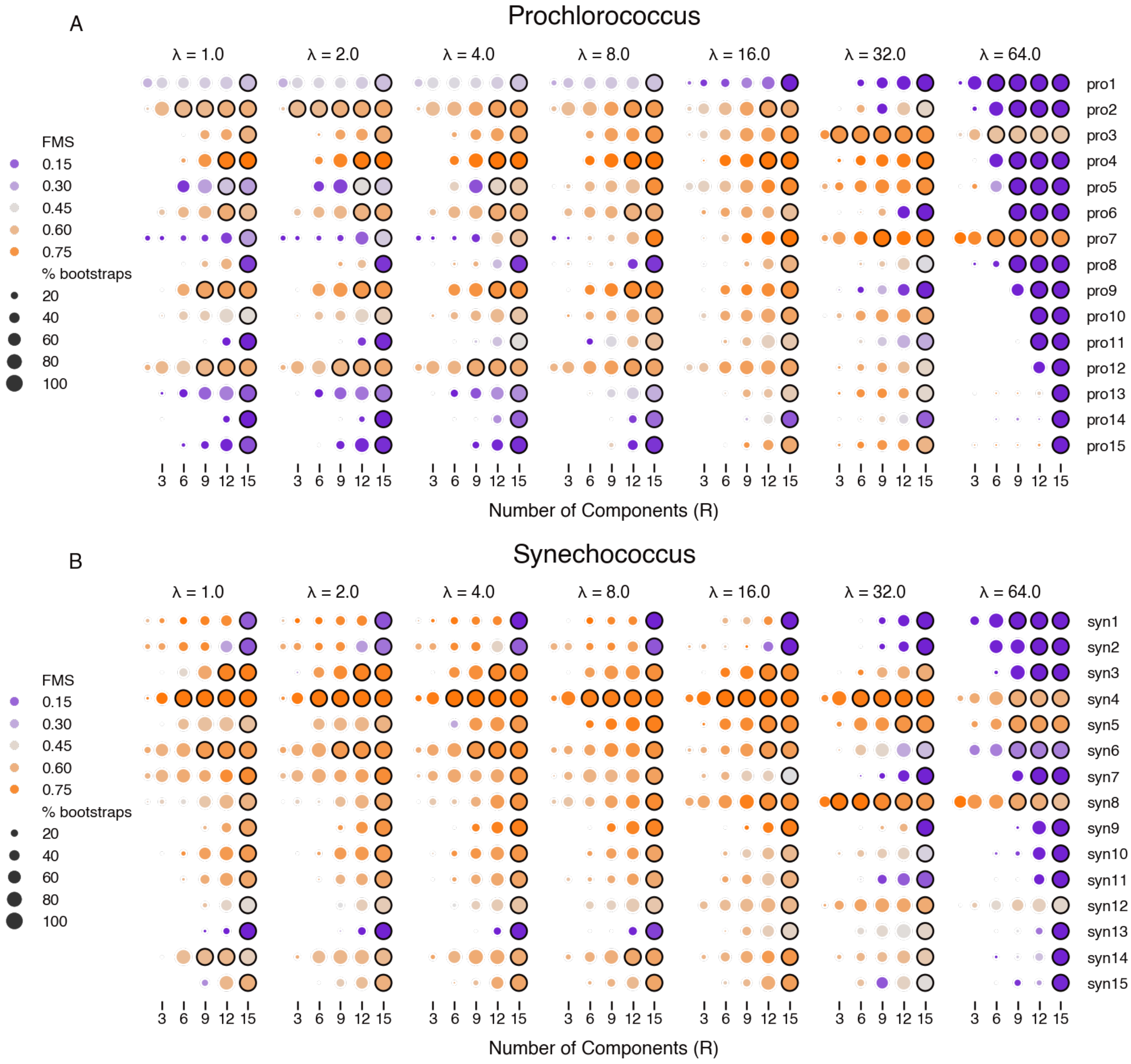


**Fig. S12. Model components are robust across an exponential range of sparsity coefficient values.**

Median cross-validated component factor match score (FMS) of (A) *Prochlorococcus* and (B) *Synechococcus* models parameterized with different numbers of components (*R*) and sparsity coefficients (λ). All models were aligned to a reference model parameterized with best fit parameters (as determined by cross-validated grid search), and FMS values were calculated between each component and its best match in the reference model. Marker color indicates median FMS of 30 bootstrapped models, marker size indicates the percentage of bootstraps in which the component was detected, and markers with a black outline indicate components that were detected in all 30 bootstraps.

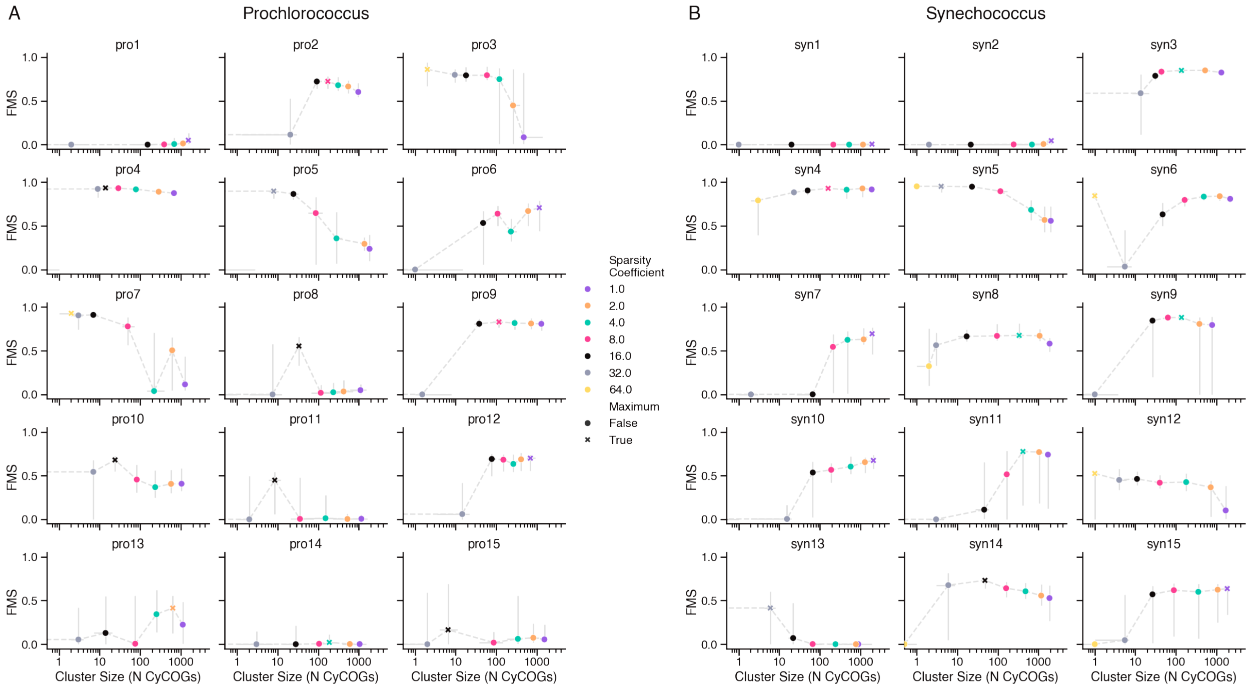


**Fig. S13. Component robustness and cluster size as a function of sparsity coefficient.** Median cross-validated component factor match score (FMS) of (A) *Prochlorococcus* and (B) *Synechococcus* models parameterized with different sparsity coefficients, indicated by marker color. The maximum median FMS observed for each component is identified by an 'x' marker. X-axis shows median cluster size, equal to the number of genes (CyCOGs) with non-zero component weights. Horizontal and vertical bars indicate inner 50th percentile among 10 model bootstraps.

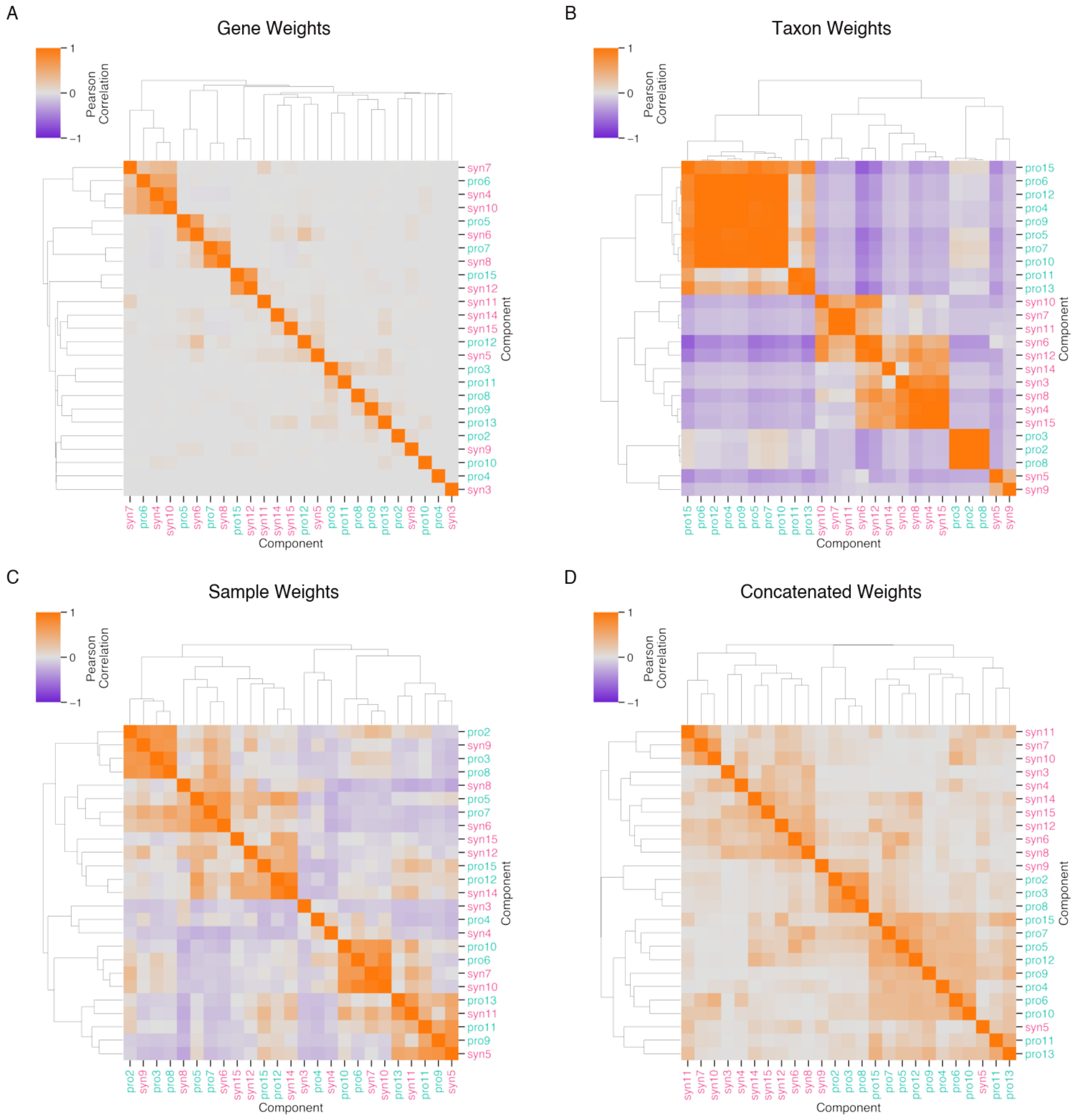


**Fig. S14. Clustered heatmaps of pairwise correlations of component weight profiles.** Hierarchically clustered heatmaps of pairwise Pearson correlations between (A) gene weight, (B) taxon weight, (C) sample weight, and (D) concatenated gene, sample and taxon weight profiles of robust *Prochlorococcus* and *Synechococcus* components.

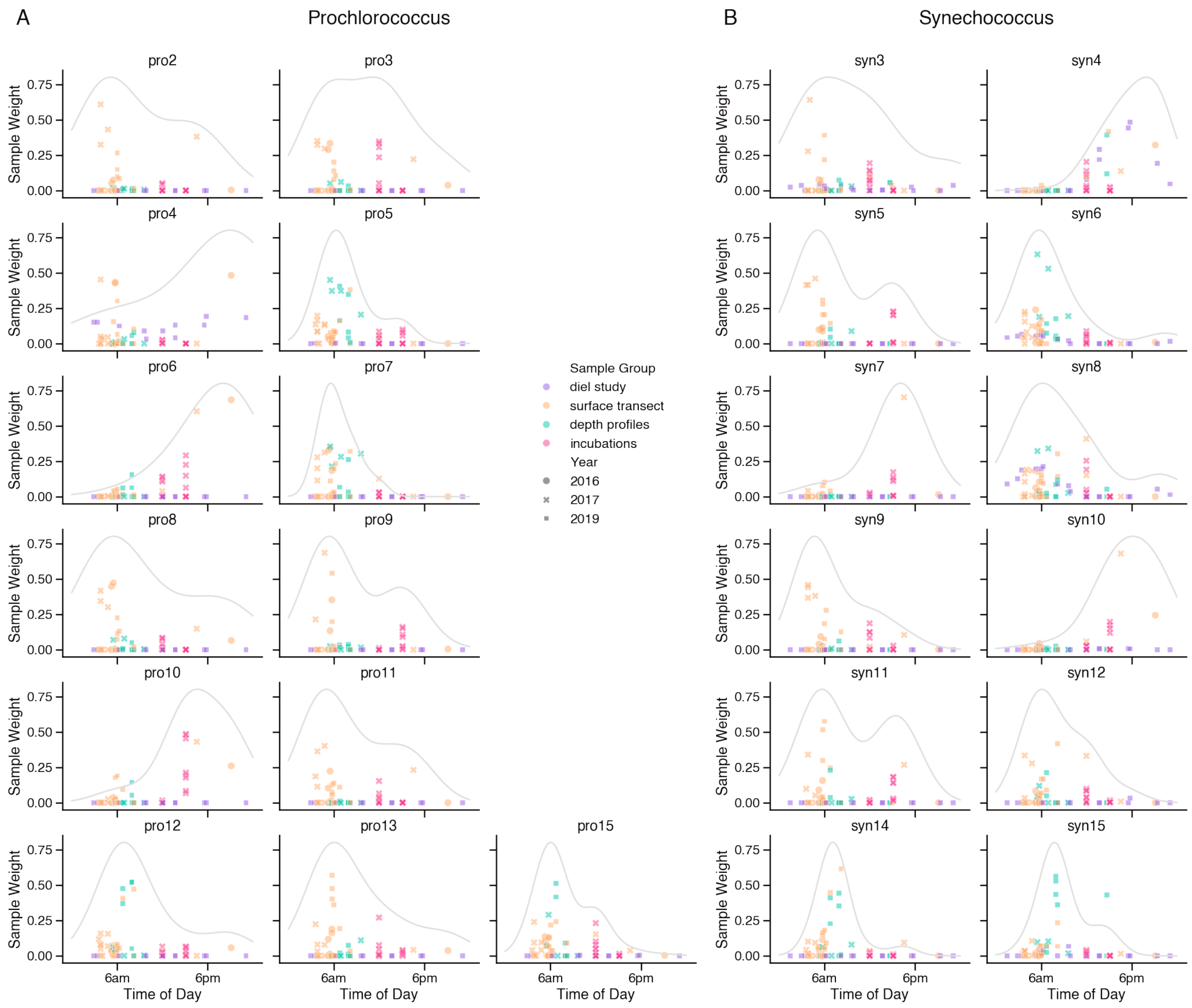


**Fig. S15. Estimation of diel peak expression time of clusters via weighted KDE.**

Sample weights of (A) *Prochlorococcus* and (B) *Synechococcus* components by time of day of sample collection. Grey lines show kernel density estimate (KDE) of sampling times, weighted by sample weight and corrected for uneven sampling, as an estimate of peak expression time of each cluster.

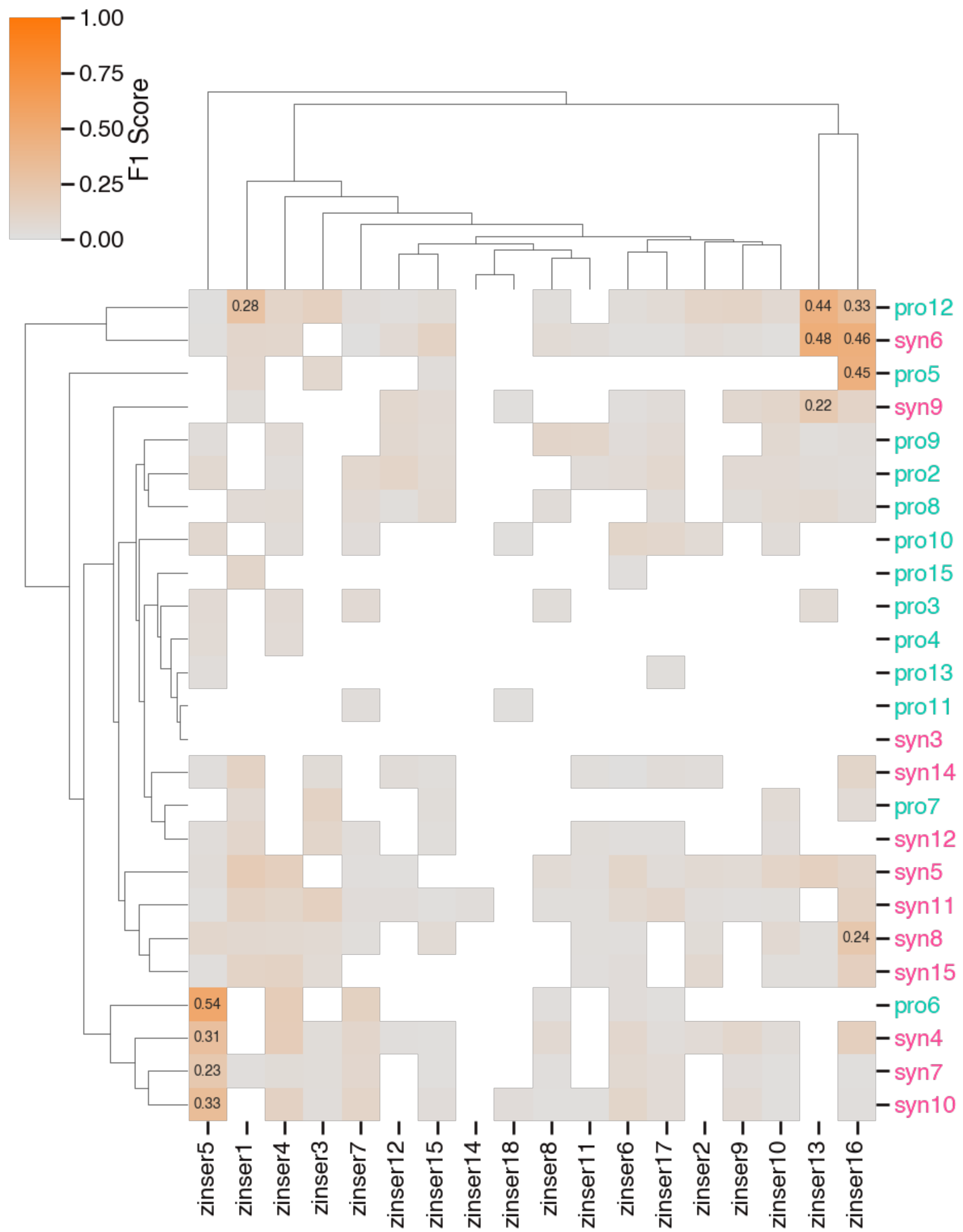


**Fig. S16. Comparison of Barnacle cluster gene membership to previously reported diel clusters of cultured *Prochlorococcus* MED4.**

Hierarchically clustered heatmap of F1 scores comparing CyCOG composition of Barnacle clusters against clusters previously derived from transcriptomes of *Prochlorococcus* MED4 grown in culture over a simulated day/night cycle (Zinser et al., 2009). Clusters without any overlapping CyCOGs are left blank and significantly similar clusters are annotated with the F1 score (adjusted $p < 0.05$).

**Legends for data S1 to S11 (separate files)**

**Data S1. *Prochlorococcus* component profiles.**

Median weight profiles for each *Prochlorococcus* component, including list of associated CyCOGs with corresponding gene weight, bootstrap support, and consensus annotation.

**Data S2. *Synechococcus* component profiles.**

Median weight profiles for each *Synechococcus* component, including list of associated CyCOGs with corresponding gene weight, bootstrap support, and consensus annotation.

**Data S3. Enrichment analysis.**

Significantly enriched KEGG pathways associated with each component, and compiled consensus annotations for each CyCOG.

**Data S4. MED4 CyCOGs.**

Mapping of *Prochlorococcus* MED4 genes to associated CyCOGs.

**Data S5. Nitrogen and iron acclimation clusters.**

CyCOGs, genes, and annotations for clusters associated with acclimation to nitrogen and iron scarcity.

**Data S6. Sample metadata.**

Metadata file detailing sampling conditions for all metatranscriptomes used in this study.

**Data S7. Genome metadata.**

Metadata file for CyCOG v6 reference genomes, including updated clade assignments.

**Data S8. CyCOG v6 database.**

Tarball of CyCOG v6 database, including reference genomes and annotation data.

**Data S9. Reference sequence phylogenies.**

Tarball of *Prochlorococcus* and *Synechococcus* reference genome phylogenies, used to update clade assignments.

**Data S10. *Prochlorococcus* transcript abundance data.**

A netCDF file of raw and normalized *Prochlorococcus* transcript abundance data, aggregated by CyCOG and organized into an `xarray.Dataset' tensor data structure.

**Data S11. *Synechococcus* transcript abundance data.**

A netCDF file of raw and normalized *Synechococcus* transcript abundance data, aggregated by CyCOG and organized into an `xarray.Dataset' tensor data structure.
